# The marine worm *Capitella teleta* is sensitive to neurochemical manipulation, as revealed via a novel high-throughput behavioural tool

**DOI:** 10.1101/2024.12.02.626409

**Authors:** Maxwell Hendershot, William Andrew Thompson, Andrea Murillo Ramos, Olivia Bradshaw, Joanna Y. Wilson

**Affiliations:** Department of Biology, McMaster University, 1280 Main St. West, Hamilton, ON L8S 4K1, Canada

**Keywords:** Pharmaceuticals and personal care products, thigmotaxis, invertebrate, locomotion, antidepressant

## Abstract

Anthropogenic impacts have led to increases in contaminants in marine habitats. The presence of compounds such as pharmaceuticals, personal care products, and pesticides is a concern, as these compounds have been shown to act as neurochemicals in aquatic organisms. However, screens of the multitude of chemicals found environmentally have yet to be carried out. *Capitella teleta* are marine annelid polychaete worms that live in the sediment of estuarian environments, acting as ecosystem engineers for marine habitats. This study aimed to develop a behaviour tool to identify neurochemical pathways involved in controlling locomotion that could be applied to screen the impacts of toxicants in the environment using this key invertebrate species at multiple life stages. Adult and juvenile life stages (2 weeks post-emergence) of *Capitella teleta* were observed in Petri dishes (adults only) or 6-well plates (adults and juveniles), and conserved behavioural responses were isolated, such as their velocity, distance travelled, the time to reach the edge of the arena, and time spent at the periphery. We exposed juvenile and adult worms to nicotine (acetylcholine agonist) as a proof of concept. We noted similar disruptions to locomotion at both life stages, with low doses of nicotine stimulating movement and higher doses reducing locomotion. From here, we exposed juvenile worms to fluoxetine (serotonin reuptake inhibitor), phenobarbital (GABA agonist), and apomorphine (dopamine agonist). The behaviour of juvenile worms can be altered by exposure to fluoxetine, phenobarbital, and apomorphine. Fluoxetine and phenobarbital exposure reduces movement at high doses, but fluoxetine influences the amount of time at the periphery of the arena. Apomorphine produced modest changes in locomotion compared to other chemicals tested. Genome searching, as well as transcriptomics, were used to confirm the presence of neurochemical pathways in *Capitella teleta*. Our results demonstrate that *Capitella teleta* is a viable model for behavioural work, that several neurochemical pathways contribute to locomotory behaviour, and that this assay may be useful as a screen for contaminants found in marine habitats.

## Introduction

Chemical contaminants, such as pharmaceuticals, are being detected at increasing levels in marine habitats (Islam and Tanaka, 2004; Prichard and Granek, 2016; Hajji and Lucas, 2024). These compounds often dissolve or get absorbed into the sediment of marine environments depending on their chemical properties, making sediments a large sink for toxicants (Chiaia-Hernández *et al*., 2022) Many species, such as invertebrates, spend their time in or on top of sediment, allowing for the uptake of a variety of contaminants through ingestion (Waring and Maher, 2005; Wright, Thompson and Galloway, 2013; Baker, Tyler and Galloway, 2014). High concentrations of contaminants within prey invertebrates can contribute to the accumulation of toxicants in higher trophic levels (Fleeger, Carman and Nisbet, 2003; Huang *et al*., 2021; Saidon *et al*., 2024). Contaminant exposures have been linked to impacts on neuronal development, endocrine signalling, maintenance of homeostasis, and the morphology of invertebrates (Koenig *et al*., 2016; Carmona, Roudeau and Ortega, 2021; Srain, Beazley and Walker, 2021), which may be suggestive of the susceptibility of these animals to toxicants. However, given the myriads of toxicants found environmentally, assessing the relative sensitivity of invertebrates to contaminants is necessary, but requires the development of high-throughput tools.

*Annelida* is a phylum of invertebrates (Ferrier, 2012) that emerged ~500 million years ago (Capa and Hutchings, 2021). *Capitella teleta* (Blake, Grassle and Eckelbarger, 2009) is a marine annelid found in the sediment of estuarine environments (Seaver, 2016). *C. teleta* are recognized as a good indicator of disturbed habitats, as they are often found near organically enriched areas, such as sewage treatment plants and oil spills (Seaver, 2016). Adults can be differentiated from juveniles by distinct physiological and morphological differences, such as the presence of sex-specific gonads (Seaver, 2016). As this animal has had its genome sequenced (Weigert and Bleidorn, 2016), this provides a genetic framework for the exploration of mechanisms of disruption by contaminants found in marine environments. Behaviour is a useful tool when investigating the effects of stressors on an animal (Anisman & Zacharko, 1986), particularly considering the role neurochemical pathways play in the regulation of behaviour (Voon *et al*., 2020). Despite this, behaviour is often an understudied effect of contaminant exposure. Alterations in behaviours could therefore resemble a change in their safety and anxiety responses, but whether contaminants can impact the behavioural profiles of an invertebrate species like C. *teleta* has not been well investigated.

Invertebrate genomes have been shown to contain conserved molecular machinery involved in the regulation of many neurochemical pathways, including serotonin (Curran and Chalasani, 2012), dopamine (Vidal-Gadea and Pierce-Shimomura, 2012), GABA (G.G. Lunt, 1991), and acetylcholine (Kim and Lee, 2018). Typically, these systems are involved in cognitive and behavioural functions (serotonin; Bacqué-cazenave et al., 2020), reward regulation, food seeking, and learning (dopamine; Barron et al., 2010; Sawin et al., 2000), attention, cellular/synaptic physiology, and motor control (acetylcholine; Colangelo et al., 2019; Mille et al., 2021) and locomotion, reproduction, and stress (GABA; Paredes & Anders, 1992). However, whether these systems exist in *C. teleta*, and are responsive to chemical agonists, has yet to be explicitly tested.

In this study, we tested the hypothesis that *C. teleta* are responsive to neurochemical manipulation as reflected by alterations to their behavioural responses. We first confirmed that neuromodulatory pathways existed within *C. teleta* using protein BLAST searching, gene ortholog alignments, and motif searches, along with the creation of phylogenetic trees for target proteins of each major system (dopamine, serotonin, acetylcholine, and GABA). Transcriptomic levels were determined for several targets within each pathway. To test whether animal performance could be manipulated by neurochemicals, we developed a high-throughput behavioural test for use in *C. teleta*. We first assessed whether adult worms could be used in a locomotory assay in petri dishes. Following which, we compared responses in juvenile and adult worms using nicotine (acetylcholine receptor agonist; Xiao et al., 2020) in a high throughput multi-well plate assay. We exposed juvenile worms to either fluoxetine (serotonin transporter inhibitor; Nutt et al., 1999), apomorphine (dopamine receptor agonist; Onyameh et al., 2021), or phenobarbital (GABA receptor; Löscher & Rogawski, 2012) at 0, 1, or 10 µM. The total distance moved (cm), time spent moving (s), time spent at the edge (s), and maximum velocity (cm/s) were measured to determine if there were any alterations in behaviour caused by chemical exposure. These results demonstrate that *C. teleta* have the molecular machinery to respond to neuroactive compounds and exhibit greater responses to serotonergic and GABAergic manipulation.

## Methods

### Culture Maintenance

Juvenile and adult *C. teleta* were obtained from a laboratory culture (Blake et al., 2009; shared by S. Hill, Michigan State University) that was maintained as previously described (Seaver, 2016). *C. teleta* were held in 4.5-inch culture dishes (Carolina Biological Supply, Burlington, USA) filled with 32 parts per thousand (ppt) artificial filtered seawater (AFSW; Instant Ocean, Spectrum Brands, Blacksburg, USA) at 15-16°C. Their food and substrate consisted of artificial marine mud made from mushroom compost (Trade Technocrats LTD, Scarborough, Canada), powdered kelp (Van Beek’s Landscape Supply, Oakville, Canada), and AFSW which were incubated at room temperature for at least 3 weeks. This allows the compost to ferment and promote the growth of bacteria, increasing the nutritional quality of the mud. Large particulates were sieved out and the mud was frozen at −20°C for 24 h minimum and aerated for 24 h before use. The artificial mud (1 Tbsp for bowls ≥8 weeks old, ½ Tbsp for bowls <8 weeks old), along with 1 TetraMin tropical flake (Tetra, Spectrum Brands, Blacksburg, USA), were used to feed the worms and provide a substrate for burrowing. A 75% water change was performed twice a week to maintain salinity and the worms were fed once a week, during one of the water changes. Bowls were sifted every 2 weeks to search for dead/sick worms, separate brood tubes, and sex the worms to remove hermaphrodites. During the sifting, *C. teleta* were placed into large petri dishes filled with AFSW. While in the large petri dishes, the worms were counted, sexually identified, and those with brood tubes were collected. Sex was identified by sexually dimorphic characteristics such as a genital spine (male) or the presence of oocytes (female), under a light microscope (Zeiss, Oberkochen, Germany). Brood tubes were placed into small Petri dishes (6 cm in diameter) filled with AFSW and kept separately from the rest of the bowl and checked daily for the emergence of larvae under a light microscope. This allowed us to prevent larvae from emerging within the culture bowl so that the age of the worms remained consistent within the bowls. The water was replaced twice a week. To maintain genetic diversity, new culture bowls were made by combining emergent animals from at least 4 brood tubes from females (but not hermaphrodites). The larvae from all brood tubes were combined in a large petri dish with AFSW and 40 larvae were randomly selected and placed into bowls with ½ Tbsp mud, AFSW, and a flake of food. The mothers were returned to their original bowls. After 8 weeks, *C. teleta* reached maturity and were sifted for the first time, sex was determined; males and females moved into a new bowl.

### Chemicals

To establish our behavioural model, we used nicotine (CAS: 65-316, 2 mM; Sigma, St. Louis, USA; an agonist of acetylcholine receptors; Xiao et al., 2020), as it has been shown to be stimulatory in *Caenorhabditis elegans* (Maupas, 1900; Feng et al., 2006; Sobkowiak et al., 2011). To assess major neurochemical pathways that could modulate behaviour, worms were exposed to fluoxetine (CAS: 56296-78-7, Sigma, St. Louis, USA; modulator of synaptic serotonin; Nutt et al., 1999), apomorphine (CAS: 41372-20-7, Cedarlane, Burlington, Ontario, Canada; an agonist of dopamine receptors; Onyameh et al., 2021), and phenobarbital (CAS: 50-06-6, Toronto Research Chemicals, Toronto, Ontario, Canada; an agonist of GABA receptors; Löscher & Rogawski, 2012). Stocks were made for each chemical (2000 µM for nicotine, 1000 µM for fluoxetine, apomorphine, and phenobarbital; stored at −20°C), and then diluted in AFSW to test concentrations for animal exposure (0, 2, and 20 µM for nicotine & 0, 1, and 10 µM for fluoxetine, apomorphine, and phenobarbital). Fluoxetine, nicotine, and phenobarbital were soluble in water; apomorphine was dissolved in DMSO and the exposure also included a 0.01% DMSO control.

### Behaviour

#### Adult behaviour

Adult worms (12-18-weeks postemergence) were sifted and individually placed inside a central circle with a radius of 1 cm (used for a consistent starting position) inside of a Petri dish (60×15 mm petri dish, Fisher Scientific, Massachusetts, USA). To examine their basal behaviours, hundreds of worms were recorded in 20–30-min intervals inside a Danio Vision behavioural platform (Noldus, Wageningen, Netherlands), which was connected to a chiller (Thermo Fisher Scientific, Waltham, Massachusetts, USA) to maintain temperature. This established their peak activity after movement to the novel arena. We observed that when adults were placed into a novel environment (the arena), there was an immediate induction of exploratory behaviour, and a propensity to move to the edge of the arena. On this basis, we used the first 10 min upon placement in a novel environment as our metric for comparing behaviour effects in adult worms. A circle was drawn at the centre of the circular Petri dish, which encompassed ~20% of the total arena to standardize the location of the worm at time zero. Adult worms were exposed to nicotine for 1 h in an incubator (held at 15°C), before being moved into Petri dishes and recorded for 10 min at 15°C in total darkness. The distance moved (cm), maximum velocity (mm/s), time to the edge of the arena (s), and time to first movement (s) were measured to assess changes in locomotion in the petri dish. A similar behavioural assessment was carried out using a 6-well plate. Given the difference in the size of the Petri dish (60 x 15 mm) compared to the 6-well plate (35 x 17 mm), periphery-seeking behaviours could not be monitored in the 6-well plate. The distance moved and maximum velocity was calculated using EthoVision XT software (version 15.0.1416; Noldus, Wageningen, Netherlands), and the videos were manually examined to determine the time required for the first movement to contact the edge of the arena. The manual scoring of videos was carried out blind to the treatment group by a single assessor.

#### Juvenile Behaviour

Juvenile worms (2 weeks post-emergence) were assessed for locomotory behaviour in a 6-well plate. This offered three advantages; juvenile worms are available earlier than adult worms, are prior to sexual maturation (thus decreasing the number of animals to test), and are a more appropriate size for measurements in a 6-well plate. The basal behaviour of the juveniles was assessed by placing worms in a 6-well plate (1 worm per well) and recording them for 3 h. From these videos, we observed that a considerable acclimation time was required, as increases in the movement of juveniles occurred 1 h after placement into the arena (SI figure 12, 13). Thus, we recorded juvenile behaviour for 10 min after 1 h of being placed in the arena. We used the acclimation period inside the arena to act as our exposure window to chemicals (1 h), matching the exposure time used in the adult studies. To validate the 6-well plate as an arena, we performed preliminary trials to observe the basal behaviour of juvenile *C. teleta* and tested the effects of nicotine on juveniles and adults within the well plate, the same chemical used in the Petri dish assay. Measures of the distance moved (cm), and maximum velocity (mm/s) of juvenile *C. teleta* were collected using the Ethovision XT software as described above. Due to the animals being in the behavioural arena for 1 h before recording, we could not record the time required for their first movement and time to reach the edge of the arena. Like adults, juvenile *C. teleta* exhibit a propensity to seek the edge of the arena, so the EthoVision XT software was used calculate the total duration that *C. teleta* juveniles spent at the edge of the arena, with the outside area consisting of a 5mm circle around the edge of the well. All trials were recorded at 15°C in total darkness.

### Identification and Selection of Neuronal Targets

The *C. teleta* genome (NCBI:txid283909) was searched using blastp (Altschul et al., 1990) in NCBI for evidence of the dopamine, serotonin, acetylcholine, and GABA pathways. The blast search was performed with gene orthologs in *C. elegans* (NCBI:txid6239) and *Homo sapiens* (Linnaeus, 1758; NCBI:txid9606). Genes from *C. elegans* were identified in WormBase (Sulson and Waterston, 1998). The specific protein (gene name, accession number) used in blast for each neuronal pathway is listed in SI Tables 1, 6, 11, & 16. Top hit putative orthologs were identified by the highest max score and lowest e-value. These gene sequences were used in gene alignments using NCBI Multiple Alignments (Papadopoulos and Agarwala, 2007) with the *C. elegans* and human ortholog sequences to assess sequence similarity by looking at the percent identity, positives, and gaps.

The *C. teleta*, *C. elegans,* and *H. sapiens* orthologs were used in InterProScan (Jones *et al*., 2014) searches for conserved motifs to confirm gene identity for the serotonin transporter and dopamine, acetylcholine, and GABA receptor sequences. Confirmation of *C. teleta* gene identity was achieved by ensuring the protein domains from the motif search matched the function of the chosen gene and stayed consistent with the motifs of the *C. elegans* and *H. sapiens* motifs. Clustal Omega (Madeira *et al*., 2024) was used to align the sequences between the 3 species, this alignment was used with the information from the InterProScan searches to highlight conserved motifs.

Phylogenetic trees were made to examine the evolutionary history of the SERT, dopamine receptor, acetylcholine receptor, and GABA-A receptor. To construct these trees, protein sequences from diverse animals were taken from NCBI (Altschul et al., 1990), including genes from mammals, reptiles, fishes, and invertebrates. A list of all organisms used and the accession number for each protein can be found in SI Tables 5, 10, 15, & 20. The protein sequences were aligned with Clustal Omega (Madeira *et al*., 2024), set with default parameters. TrimAI (v. 1.3; (Capella-Gutiérrez, Silla-Martínez and Gabaldón, 2009) was used with the automated 1 setting to mask the sequences, accessed through the Phylemon2 web server (v. 2.0; Sánchez et al., 2011). ModelTest (Flouri *et al*., 2015; Darriba *et al*., 2020) was used to identify the best parameters for each protein alignment and used in RaxML (v8.2.12, Stamatakis, 2014), to generate phylogenetic trees with bootstrapping (100 replicates). Phylogenetic analyses for the SERT and acetylcholine receptor used the LG4X+I model of amino acid substitution, while the dopamine and GABA receptors used the, LG+I+G and LG+4G models of amino acid substitution. The final trees were viewed using iTol (v6, Letunic & Bork, 2024).

### RNA Extraction, Sequencing, and Differential Gene Expression of Target Genes

Worms were placed on 0.6% cornmeal agar plates (Sigma Aldrich, St. Louis, USA) made with filtered seawater for 4 h to clear gut content. Juveniles were pooled with 10 worms per biological replicate and all adult sexes were pooled with 5 worms per sex per biological replicate, and worms were snap frozen and stored at −80C. All groups had a total of 3 biological replicates. Total RNA was extracted from each biological replicate using the TRIzol extraction (Thermo-Fisher, Massachusetts, USA) method combined with RNeasy® Plus Mini kit (Qiagen, Hilden, Germany) as performed by (Gayral et al., (2011). Total RNA samples were sent for library construction and sequencing to the McMaster Genomics Facility (Hamilton, Canada). RNA quality and integrity were evaluated for all samples using a 2100 BioAnalyzer (Agilent Technologies, California, USA) before library preparation. Due to the nature of the RNA extraction, only one peak was expected rather than two as the 28s rRNA peak is denatured and migrates to the 18s rRNA peak (Gayral *et al*., 2011). As a RIN value is not possible without both 28s and 18s rRNA peaks, RNA quality was inspected visually, for a single peak and the absence of peaks at smaller fragments of rRNA that would suggest degradation. mRNA was enriched using the NEBNext® Poly(A) mRNA Magnetic Isolation Module per the manufacturer’s instructions (New England Biolabs, Massachusetts, USA). RNAseq libraries were made from the enriched mRNA using the NEBNext® Ultra II Directional RNA Library Prep Kit for Illumina per the manufacturer’s instructions (New England Biolabs, Massachusetts, USA). After library preparation, their quality, size, and concentration were determined for equal input into the sequencer. High throughput sequencing was performed using the Illumina NextSeq 2000 platform (Illumina, CA, USA) with a P2 100 cycle flow cell in a setup of paired, 2×50 base pairs.

The quality of the reads was checked before alignment using FastQC (version 0.12.0). Reads that were low quality and adapters were trimmed using Trimmomatic (version 0.38) with sliding window parameters set to 4 base pairs with an average minimum Phred score of 15 (Bolger, Lohse and Usadel, 2014). Reads with a minimum length of 36 base pairs were kept (Bolger, Lohse and Usadel, 2014). The trimmed reads were aligned to the *C.teleta* genome (GCA_000328365.1 from the Ensemble Metazoa Database Release 52; Simakov et al., 2013) using HISAT2 (version 2.2.1; D. Kim et al., 2019). Alignments were evaluated using FastQC (version 0.12.0). Gene expression estimates were computed using Stringtie (version 2.2.0) with settings for first-stranded synthesis and the reference annotation file (Capitella_teleta_v1.0.52.gtf.gz from the Ensemble Metazoa Database Release 52) to determine the number of reads per gene. Differential expression analyses were conducted using the read count output from Stringtie into DESeq2 (v 1.42.0) in R (v 4.3.2). Before running the analysis, read counts below 10 were filtered out. A likelihood ratio (LRT) hypothesis test was run using the *DESeq* function with a full design model including all sex conditions compared to a reduced model of 1. The conditions of sex included juveniles, males, and females. Gene expression plots were made using RStudio (v 2024-06-14, Posit Team, 2024), using the plotcounts function to plot out the objects made by the analyses.

### Statistics

The behavioural data are presented as untransformed means ± SEM, but a square root transformation was performed in all cases where parametric tests were used. For the total distance travelled and maximum velocity in the Petri dish assay, a one-way analysis of variance (ANOVAs) followed by Tukey’s multiple comparison test was used to determine statistical differences among groups. For the time to first movement and time to reach the arena’s edge, a non-parametric Kruskal-Wallis test was performed. Behavioural statistical analyses were performed in Prism (v8.0). For all 6-well plate assays, a non-parametric Kruskal-Wallis test was performed for the distance moved, maximum velocity, and time spent at the edge because the data could not meet the assumptions of homoscedasticity and normal distribution following transformations. A Dunn’s multiple comparison test was performed if any main effects were significant. Behavioural statistical analyses were performed in R studio (v 2024-06-14, Posit Team, 2024). Alpha was set to 0.05 in all statistical tests and different letters denote statistical differences across groups.

## Results

### Major neurochemical pathways are present in C. teleta

Orthologs of the serotonin transporter and acetylcholine, dopamine, and GABA receptors were present in the *C. teleta* genome (SI Table 1, 6, 11, 15), as were other key genes in these pathways (SI Table 1, 2, 6,7, 11, 12, 16, 17). For the primary gene targets for the chemical manipulations, the *C. teleta* genes had similar motifs as the human and *C. elegans* orthologs (SI Figure 2, 5, 8, 11). Phylogenetic analyses of these gene targets placed the *C. teleta* genes with taxonomically related species with reasonable bootstrap support. All of the gene targets, and other key genes in these pathways were expressed in both adult (male and female) and juvenile worms (SI Figure 2, 4, 7, 10). This suggests that the acetylcholine, dopamine, GABA, and serotonin pathways are present in both juvenile and adult *C. teleta*.

### Nicotine exposure alters C. teleta locomotion

Adult *C. teleta* worms increased their total distance moved (Figure 1A; p=0.0322) and maximum velocity (Figure 1B; p=0.0422) after exposure to 2 µM of nicotine in a petri dish arena. However, after exposure to 20 µM, they decreased their total distance moved (Figure 1A; p=0.0075), but not their maximum velocity (Figure 1B). Adult *C. teleta* took longer to reach the arena’s edge after exposure to 20 µM of nicotine (Figure 1C; p= 0.00060), and their first movement was delayed compared to other treatment groups (Figure 1D; p=0.0025).

**Figure 1.**
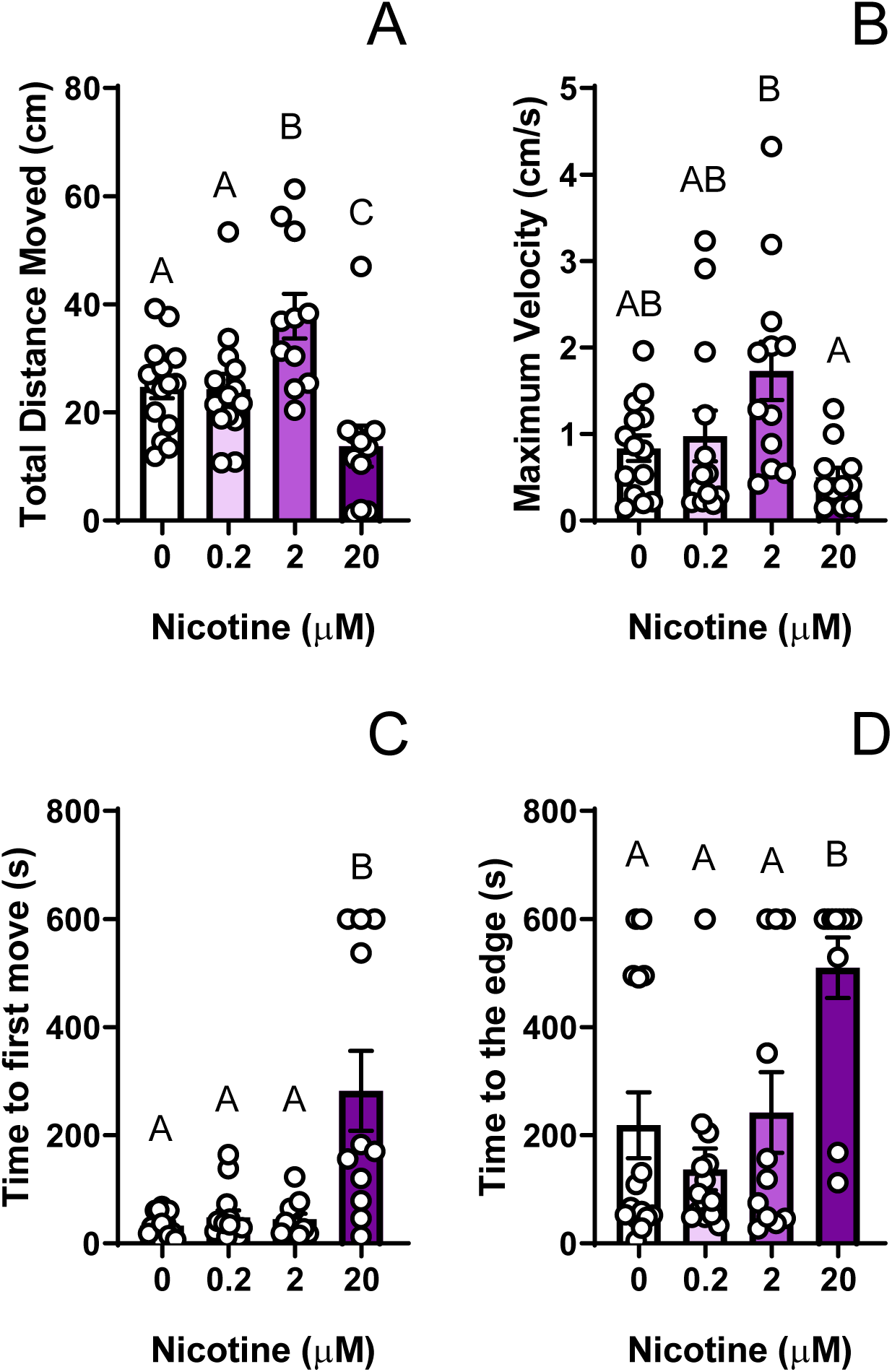
Nicotine exposure modulates *C. teleta* adult locomotory behaviour in a Petri dish arena. Adult *C. teleta* worms (n=11-14 per treatment) were exposed to a seawater control or 0.2, 2, or 20 µM nicotine for 1 h in darkness at 15°C. After exposure, animals were moved to a novel arena (petri dish) to induce exploratory behaviour. Worms were video recorded in darkness at 15°C for 10 minutes and (A) total distance moved, (B) maximum velocity, (C) time to reach the arena’s edge, and (D) time to first movement were determined. Data are presented as means ± SEM, different letters denote statistical differences across groups.

Nicotine exposure altered the total distance moved in adult *C. teleta* in the smaller arena of a 6-well plate. The distance moved was less (p=0.0016; Dunn’s Test; Figure 2A) for worms exposed to 20, but not 2 µM of nicotine relative to control. The distance moved at 20 µM was also lower than the 2 µM exposure (p=0.0040; Dunn’s test; Figure 2A). The time spent at the edge of the arena was decreased by nicotine exposure; worms exposed to 20 µM were different from controls (p=0.034; Dunn’s Test; Figure 2B). There was no difference between 2 µM and control, or between 2 µM and 20 µM treatment groups. Similar to distance moved, nicotine exposure decreased the maximum velocity of worms exposed to 20 µM relative to the control (p=0.0040; Dunn’s Test; Figure 2C) and 2 µM (p=0.0016; Dunn’s Test; Figure 2C); there was no difference between control and 2 µM groups.

**Figure 2:**
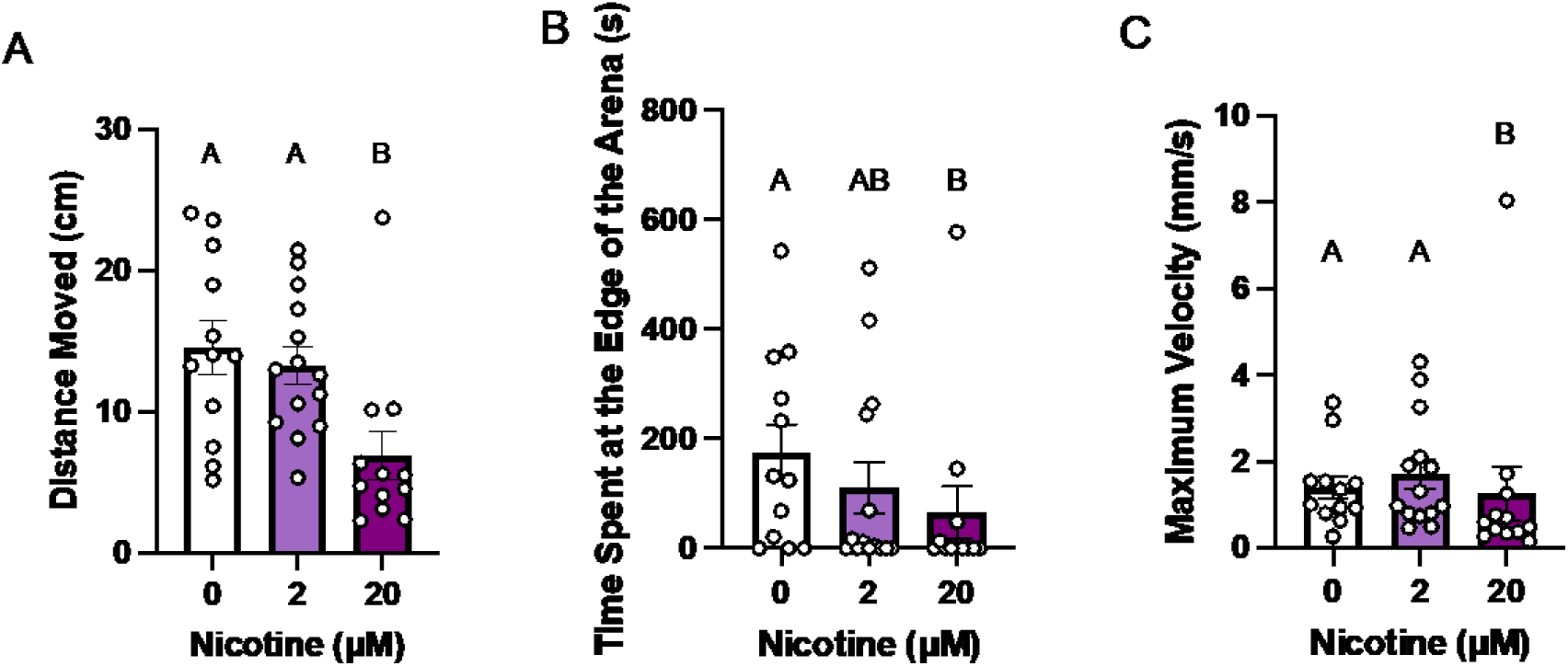
Nicotine exposure decreases *C. teleta* adult locomotion in a 6-well plate. Adult *C. teleta* (n=12-14 per treatment) were exposed to control (artificial filtered seawater), 2, or 20 µM nicotine for 1 h in darkness at 15°C. After exposure, animals were moved to a novel arena (single well from a 6-well plate) to induce exploratory behaviour. Worms were video recorded in darkness at 15°C for 10 minutes, and (A) total distance moved, (B) time spent at the edge of the arena, and (C) maximum velocity were measured. Data are presented as means ± SEM. Different letters denote statistical differences across groups.

### Nicotine exposure alters the locomotion of juvenile C. teleta

Exposure to nicotine had significant effects on the locomotion of juvenile *C. teleta* in a 6-well plate. Nicotine exposure altered the total distance moved and velocity of juvenile *C. teleta*, but not the time spent at the edge of the arena (Figure 3B). The worms increased their total distance moved (p=0.047; Dunn’s Test; Figure 3A) after exposure to 2 µM of nicotine but decreased their total distance moved (p=0.063; Dunn’s Test; Figure 3A) at 20 µM relative to the control. This resulted in a decrease in the distance moved from 2 µM to 20 µM (p<0.001; Dunn’s Test; Figure 3A). Juvenile worms exposed to 2 µM of nicotine had an increased maximum velocity compared to control (p=0.010; Dunn’s Test; Figure 3C) and 20 µM (p<0.001; Dunn’s Test; Figure 3C).

**Figure 3:**
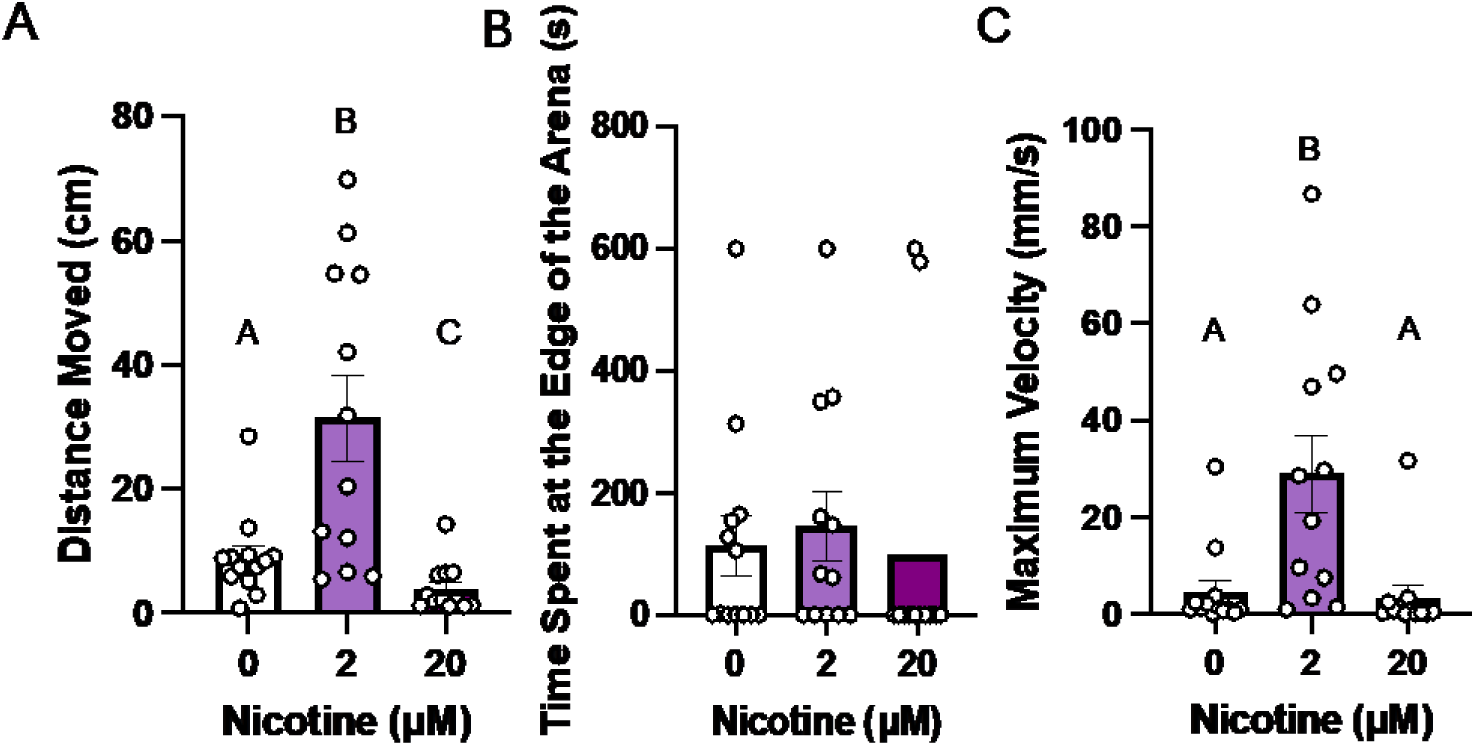
Nicotine exposure affects *C. teleta* juvenile locomotion. Juvenile *C. teleta* worms (n=12-13 per treatment) were exposed to control (artificial filtered seawater), 2, or 20 µM nicotine for 1 h in darkness at 15°C. After exposure, the worms were video recorded in darkness at 15°C for 30 minutes and (A) total distance moved, (B) time spent at the edge of the arena, and (C) maximum velocity were measured. Data are presented as means ± SEM Different letters denote statistical differences across groups.

### Fluoxetine exposure has some effect on the locomotion of juvenile C. teleta

After exposure to 1 µM of fluoxetine, the worms increased the distance moved relative to the 10 µM exposure (p=0.0272; Dunn’s Test; Figure 4A) but there were no significant differences between the 1 µM and control groups, nor the 10 µM and control groups. Exposure to 1 µM fluoxetine increased the time spent at the edge relative to the control (p=0.0052; Dunn’s test; Figure 4B). Exposure to fluoxetine did not affect the maximum velocity of the juvenile *C. teleta* (Figure 4C).

**Figure 4:**
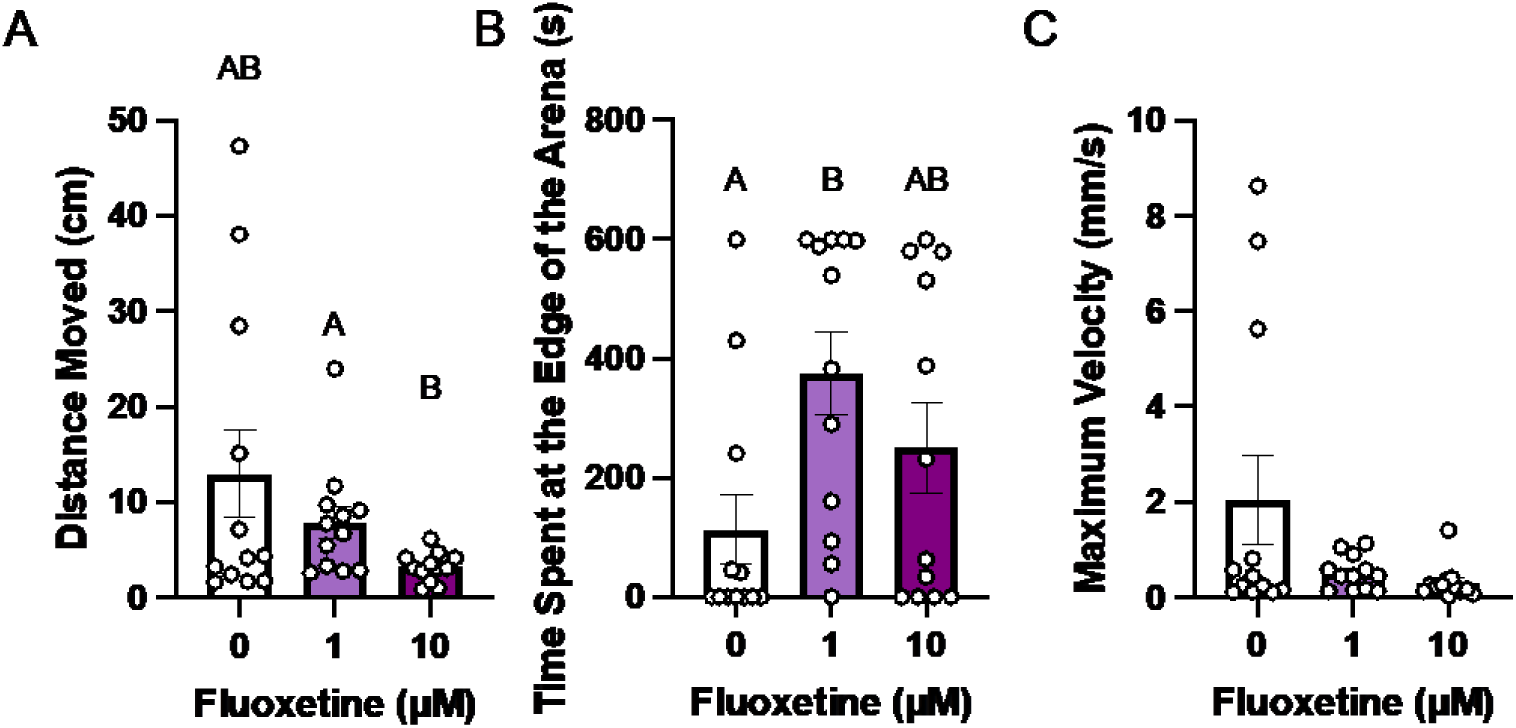
Fluoxetine exposure affects *C. teleta* juvenile locomotion. Juvenile *C. teleta* worms (n=12 per treatment) were exposed to control (artificial filtered seawater), 1, or 10 µM fluoxetine for 1 h in darkness at 15°C. After exposure, the worms were video recorded in darkness at 15°C for 30 minutes and (A) total distance moved, (B) time spent at the edge of the arena, and (C) maximum velocity were measured. Data are presented as means ± SEM Different letters denote statistical differences across groups.

### Exposure to apomorphine has a limited effect on the locomotion of juvenile C. teleta

There were no differences across treatment groups for the time spent at the edge of the arena or maximum velocity (Figure 5B & C) for juvenile *C. teleta* exposed to apomorphine. The worms increased their distance moved after exposure to 1 µM apomorphine compared to the 10 µM worms (p=0.0348; Dunn’s Test; Figure 5A), but not the control or DMSO solvent control.

**Figure 5:**
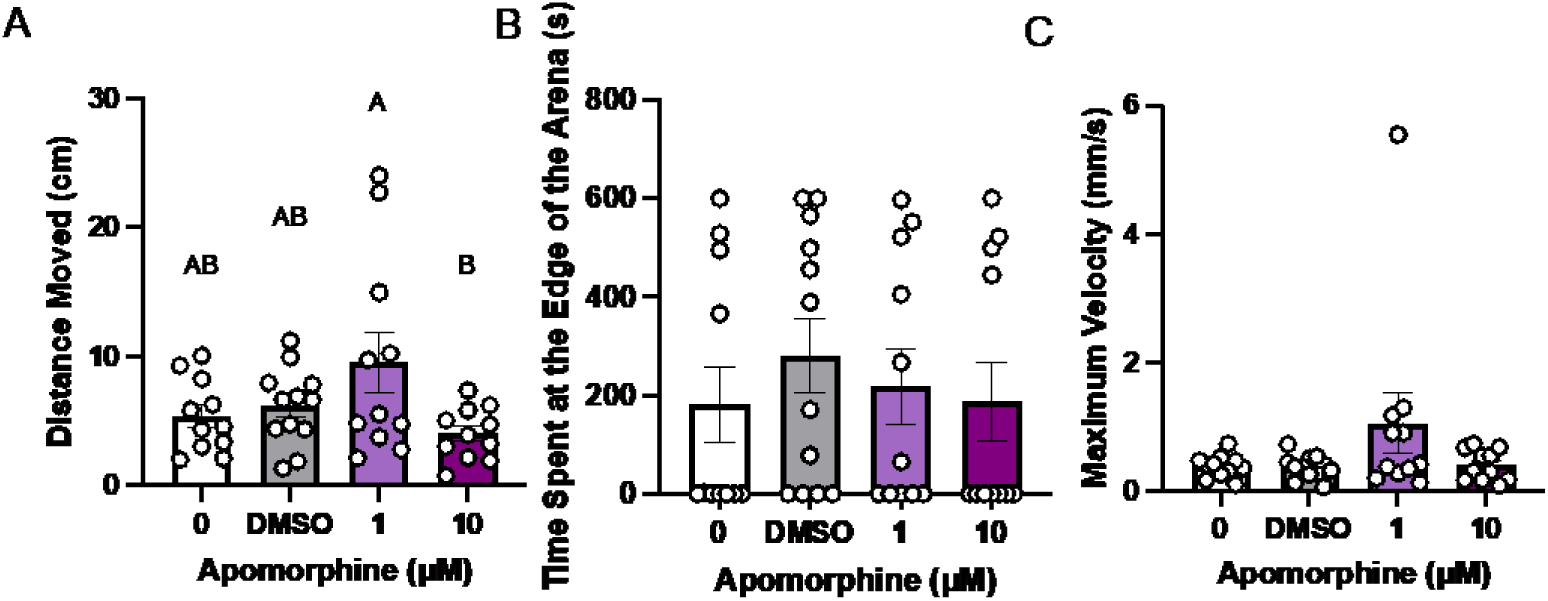
Apomorphine exposure has limited effect on *C. teleta* juvenile locomotion. Juvenile *C. teleta* worms (n=11-12 per treatment) were exposed to an artificial filtered seawater control, 0.01% DMSO control, 1, or 10 µM apomorphine for 1 h in darkness at 15°C. After exposure, the worms were video recorded in darkness at 15°C for 30 minutes and (A) total distance moved, (B) time spent at the edge of the arena, and (C) maximum velocity were measured. Data are presented as means ± SEM Different letters denote statistical differences across groups.

### Exposure to phenobarbital has inhibitory effects on the locomotion of juvenile C. teleta

Exposure to phenobarbital inhibited the locomotion of juvenile *C. teleta.* The worms decreased their distance moved after exposure to 10 µM relative to control (p=0.035; Dunn’s Test; Figure 6A) and 1 µM (p=0.022; Dunn’s Test; Figure 6A). They also decreased their maximum velocity after exposure to 10 µM compared to control (p=0.024; Dunn’s test; Figure 6C), but not to 1 µM. There were no significant differences in the time spent at the edge of the arena.

**Figure 6:**
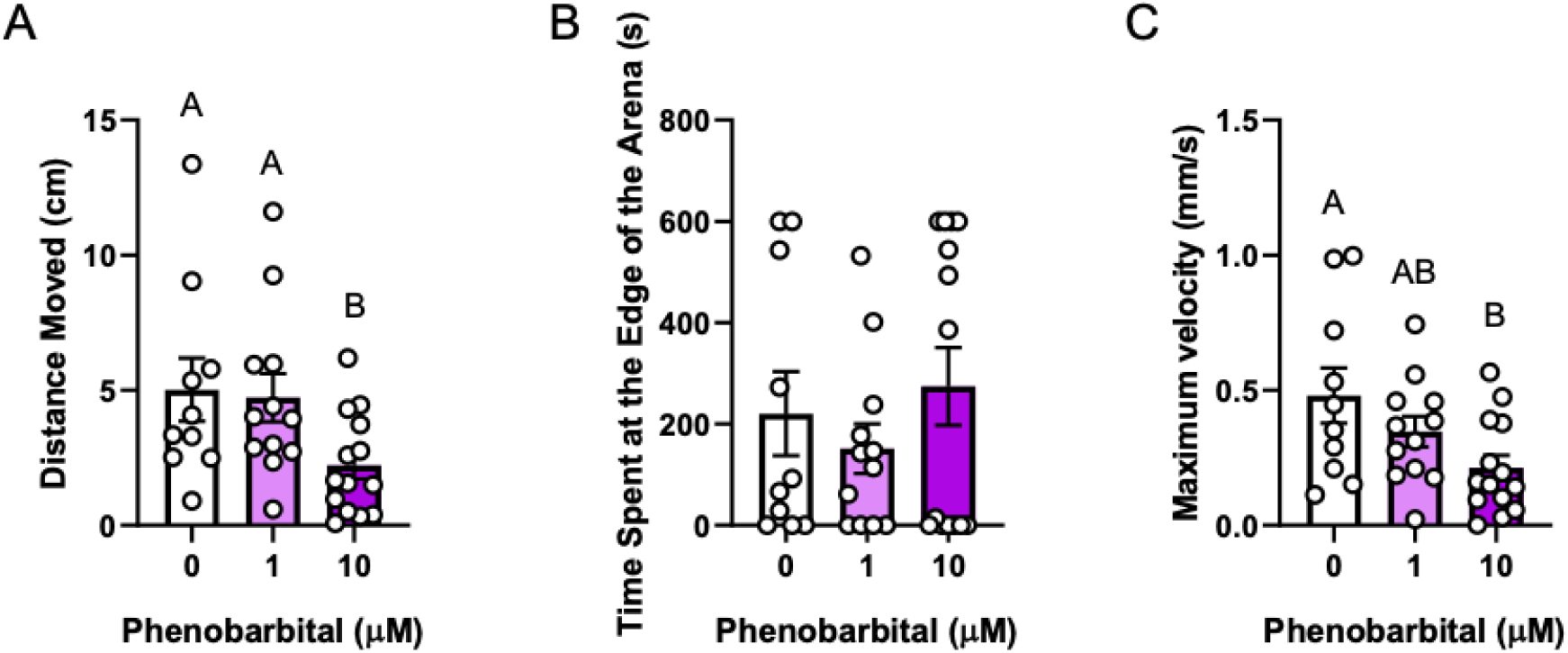
Phenobarbital has inhibitory effects on *C. teleta* juvenile locomotion. Juvenile *C. teleta* worms (n=11-14 per treatment) were exposed to an artificial filtered seawater control, 1, or 10 µM phenobarbital for 1 h in darkness at 15 °C. After exposure, the worms were video recorded in darkness at 15 °C for 30 minutes and (A) total distance moved, (B) time spent at the edge of the arena, and (C) maximum velocity were measured. Data are presented as means ± SEM Different letters denote statistical differences across groups.

## Discussion

As human-made chemical contaminants continue to rise in marine environments (Islam and Tanaka, 2004; Prichard and Granek, 2016; Hajji and Lucas, 2024) and affect species of all trophic levels (Carbery, O’Connor and Palanisami, 2018; Gilmour *et al*., 2019; Schaap *et al*., 2023), there is an urgent need to identify chemicals that may be detrimental to ecosystem health. Invertebrates are a critical group of animals, responsible for several ecosystem services, such as nutrient cycling, the decomposition of organic matter, and acting as a food source for predators (Covich, Palmer and Crowl, 1999). Due to their interaction with sediment (Waring and Maher, 2005; Wright, Thompson and Galloway, 2013; Baker, Tyler and Galloway, 2014), which serves as a major sink for chemical absorption in aquatic environments (Chiaia-Hernández *et al*., 2022), *C. teleta* represent a model organism for testing the effects of contaminant exposure. Using *C. teleta*, ecosystem engineers who stimulate environments through bioturbation (Jang *et al*., 2021), we describe a high-throughput behavioural system capable of discerning the relative sensitivity of this critical marine annelid to chemical exposure. Our results demonstrate that nicotine can act as a stimulant or a suppressor of behaviour (based on dosage), with inhibitory effects also observed with chemicals that manipulate GABA and serotonin, but with limited impacts following exposure to dopamine agonists. While most of our behavioural tests revealed alterations to the movement of *C. teleta*, we also show that compounds that modulate serotonin may influence edge-seeking behaviours, potentially pointing to conserved anxiety-like responses existing in these animals. Underpinning these results, our molecular investigation shows that the machinery for the glutamatergic, serotonergic, dopaminergic, and nicotinic cholinergic systems exist in this species. Together, these results offer insights into the neurochemical manipulation of the behaviour of this critical invertebrate species, allowing us to propose the utility of our high throughput behavioural screen to be used for testing contaminants of concern found environmentally.

The behavioural impacts of the compounds tested were tied to the mechanism of action and dosage. Nicotine interacts with the acetylcholine pathway by binding to nicotinic acetylcholine receptors (Xiao *et al*., 2020), and our results demonstrate that this neurotransmitter is involved in mediating exploratory behaviour and escape responses in *C. teleta*, with different effects depending on the concentration the worms were exposed to. Acetylcholine acts as a neurotransmitter that increases axon excitability and decreases the response threshold in some neurons (Hideki Kawai, Ronit Lazar and Raju Metherate, 2007), which could account for the stimulatory response seen at 2 µM. In contrast, exposure to a higher dosage led to an inhibitory effect, implying that the level of receptor activation may dictate the resulting phenotype. This is supported by Polli and colleagues (2015), where differences in nicotine dose had different effects on locomotory speeds in *C. elegans.* A similar result is seen with fluoxetine, which also caused inhibitory effects at the higher dosage (10 µM) exposure compared to the 1 µM exposure. Interestingly, fluoxetine increased anxiety-like behaviour at 1 µM, implying that serotonin may be involved in regulating how *C. teleta* move and orient themselves in an environment. The activation of serotonergic receptors leads to an inhibitory effect on locomotion, a result seen in both mice (Correia *et al*., 2017) and *C. elegans* (Dag *et al*., 2023). The increase in thigmotaxis (a behaviour described as when an animal “hugs the wall”; Schnörr et al., 2012) after exposure to 1 µM of fluoxetine is surprising, as thigmotaxis is linked to anxiety-related behaviours (Schnörr *et al*., 2012; Seibenhener and Wooten, 2015), and SSRIs are effective in treating anxiety disorders, helping to *reduce* anxiety (Cassano et al., 2002). This supposed increase in anxiety-like behaviour after exposure to fluoxetine is not unheard of, as fluoxetine has anxiogenic-like effects after acute exposure in rats (Liu *et al*., 2010). While our results demonstrate that the acetylcholinergic and serotonergic systems can manipulate behavioural responses in *C. teleta*, the specific receptor subtypes involved in these processes remain to be determined.

In contrast, apomorphine, a potent agonist of dopaminergic receptors (Onyameh *et al*., 2021) had little effect on *C. teleta* behaviour. This was surprising, as activation of the dopamine pathway can act as a behavioural switch from fast to slow motor patterns (Vidal-Gadea and Pierce-Shimomura, 2012), and previous studies have shown that dopamine reduces crawling speed in *C. elegans* (Sawin, Ranganathan and Horvitz, 2000). The varied response observed in the present study in response to apomorphine may be driven by our experimental paradigm. For example, dopamine has been shown to stimulate exploratory behaviour in *C. elegans* when presented with a stimulus (Hills, Brockie and Maricq, 2004), a methodology not employed here, as we solely investigated behavioural patterns following transfer. While the results here show that apomorphine can produce modest changes in behaviour in *C. teleta*, a more expansive behavioural battery testing the worm in response to stimuli may be required to fully discern the effects of dopaminergic agonism.

The GABA pathway, activated with exposure to phenobarbital, also acts as a neurochemical pathway that regulates locomotion but did not show the dose-dependent effect seen with fluoxetine and nicotine. Phenobarbital had an inhibitory effect on juvenile *C. teleta* locomotion, a result only seen at the highest dosage (10 μM) tested. Phenobarbital acts as a GABA receptor agonist (Löscher and Rogawski, 2012), and the GABA signalling pathway is considered inhibitory (Olsen, 1991). This may, in part, explain the decrease in activity noted here. GABA has been noted to be involved in anxiety disorders, and application of levetiracetam, another GABA agonist, resulted in decreased levels of glutamine levels in the thalamus, where high levels of this amino acid have been correlated with high levels of social anxiety disorder (D’costa and Shepherd, 2014). While we have generated some evidence of GABA being involved in mediating behavioural responses, measures of amino acids may be useful in determining the mechanisms of disruption of this class of compounds.

An important observation of this work was the effect arena size had on the adult stage of this animal. The validation assay in Petri dishes showed that nicotine could either stimulate or inhibit the locomotion of adult *C. teleta*, a result we were unable to recapitulate in the 6-well plates. We speculate this is due to the large decrease in arena size in a 6-well plate compared to the size of the petri dish. The worms may not have “space to behave” (Kohler, Parker and Ford, 2018), as a study by Scharf et al. (2024) found that the size of an arena affected the distance, time at the edge of the arena, and movement directionality of beetles. Likewise, the mean velocity and activity of crustaceans have been affected by arena size (Kohler, Parker and Ford, 2018). Our results further support the idea that an appropriate arena size is needed to properly assess the behavioural sensitivity of animals to contaminants, as the usage of adults in a 6-well plate would have precluded the observation of a stimulatory effect of nicotine at this life stage. Additionally, this study supports the usage of Petri dishes as an appropriate arena for the measurement of behaviour in solitary adult *C. teleta*, with the 6-well plate sufficient for the characterization of juvenile behaviour. This is an important consideration for the adaptation of this methodology for usage in other invertebrates.

Behaviour can serve as an important endpoint for toxicological testing, serving as a marker for locomotory, social, morphological disruption, or neurotoxicity in other species (Carmona et al., 2021; Diaz-Camal et al., 2022; Jones & Miller, 2008; Koenig et al., 2016; Thompson and Vijayan, 2022). The finding of this vital invertebrate species being sensitive to neurochemical manipulation allows us to further explore *C. teleta* in a toxicological setting, giving us a new high throughput, low-effort assay for use in assessing marine toxicity. Future studies should include testing for the effects of compounds at environmentally relevant concentrations, to see if *C. teleta* are sensitive to low concentrations of environmental pollutants, as well as chronic exposures starting at the larval life stage assessing longer-term exposures or the consequences of acute development exposure. This assay can also be used as a tool to investigate the different mechanisms that govern behaviour in *C. teleta* and is flexible enough to be used in different scenarios, including brood tube-related behaviours, breeding, and courtship. Adults can be used to examine sex-specific behaviours, particularly in response to compounds that target hormonal function. Overall, we have demonstrated a widely available behavioural system can be used to assess invertebrate locomotion reliably, providing a critical tool for the identification of chemicals of concern for risk management.

## Supporting information

Supplemental information for Hendershot et al 2024

Raw data for Hendershot et al 2024

## Acknowledgements

The authors would like to thank Jonathan Rahman for help with the polychaete culture, Lisa Laframboise for help with polychaete culture maintenance and adult nicotinic Petri dish exposures, and Mellissa Easwaramoorthy for support with the phylogenetic analyses.

## Author Contributions

Conceptualization (AMR, JW, MH, WAT), Data Curation (AMR, MH, WAT), Formal Analysis (AMR, MH, WAT), Funding (JW), Investigation (AMR, MH, OB), Methodology (AMR, MH, WAT), Project Admin (AMR, JW, MH), Software (AMR, MH, WAT), Resources (JW), Supervision (JW), Validation (AMR, MH), Visualization (MH, WAT), Writing (Original) (AMR, MH, WAT), Writing (Review) (AMR, JW, MH, OB, WAT)

## Data Availability

Data generated or analyzed during this study are publicly available in the Federal Research Data Repository, https://doi.org/10.20383/103.01128

