## Supplemental information for Hendershot et al 2024 for "The marine worm *Capitella teleta* is sensitive to neurochemical manipulation, as revealed via a novel high-throughput behavioural tool"

Running Title: A novel high-throughput model for toxicological testing using *Capitella teleta*

Key Words: polychaete, acetylcholine, dopamine, serotonin, GABA, locomotion

Contents: Results for the genomic searches, transcriptomics, and basal behaviour of *C. teleta*

***Identification and Selection of Targets***

Dopamine

Using *C. elegans* and *H. sapiens* dopamine receptor, tyrosine hydroxylase, dopamine ß-hydroxylase, catechol-O-methyltransferase, monoamine oxidase B, aromatic acid decarboxylase, and aldehyde dehydrogenase sequences, we found clear homologs in *C. teleta* (all accession numbers can be found in SI Table 1). The percent identities and E-values of the BLAST searches can be found in SI Table 2. All *C. teleta* sequences were found using the *C. elegans* sequences, except for Catechol-O-Methyltransferase and Monoamine Oxidase B, which were found using *H. sapiens* sequences. An alignment between *C. teleta* and *C. elegans* for Catechol-O-Methyltransferase was not possible, as the sequence for *C. elegans* was unable to be found (SI Table 2). Except for Catechol-O-Methyltransferase, all sequences had high query cover percentages and max scores (SI Table 2). The percent identity of all sequences was moderate, ranging between 27.34-66.46, and all had low e-values, with Catechol-O-Methyltransferase being the highest (SI Table 2). The sequence alignment from the Multiple Alignment aligned well, often with the *H. sapiens* sequences having higher percent identities and positives, and lower gaps, which can be seen in SI Table 3.

**SI Table 1: Accessions for proteins involved in the dopamine pathway from *Capitella teleta, Caenorhabditis elegans,* and *Homo sapiens****.* Accessions were taken off of NCBI.

| **Protein** | **Capitella teleta** | **Caenorhabditis elegans** | **Homo sapiens** |
| --- | --- | --- | --- |
| Dopamine Receptor | ELT96301.1 | NP_001024569.1 | P14416.2 |
| Tyrosine Hydroxylase | ELU11498.1 | P90986.4 | KAI4069455.1 |
| Dopamine ß-Hydroxylase | ELU08648.1 | Q9XTQ6.3 | AAH17174.1 |
| Catechol-O-Methyltransferase | ELT93114.1 | − | P21964.2 |
| Monoamine Oxidase B | ELT97776.1 | NP_001369790.1 | P27338.3 |
| Aromatic Acid Decarboxylase | ELU12210.1 | CCD63121.1 | CAG33005.1 |
| Aldehyde Dehydrogenase | ELU01961.1 | NP_498081.2 | P00352.2 |

**SI Table 2: BLAST outputs from *Capitella teleta* for proteins involved in the dopammine pathway.** BLASTS were run against *Caenorhabditis elegans* through NCBI blastp. * indicates the BLAST was run between *Capitella teleta* and *Homo sapiens.* Query Cover provides an indication of the length of each subject refers to the length of the target sequence compared to the reference sequence. Percent identity refers to the percentage of the amino acids that are identical between the sequences. Max score refers to the highest score calculated for the alignment between the sequences. The E-value refers to the number of alignments expected by chance with the calculated score.

| **Sequence** | **Query Cover (%)** | **Percent Identity** | **Max Score** | **E-value** |
| --- | --- | --- | --- | --- |
| *Dopamine Receptor* | 89 | 33.73 | 199 | 8e^-60^ |
| *Tyrosine Hydroxylase* | 77 | 41.67 | 343 | 2e^-113^ |
| *Dopamine β-Hydroxylase* | 79 | 37.15 | 362 | 1e^-116^ |
| *Catechol-O-Methyltransferase** | 47 | 27.34 | 54.3 | 1e^-09^ |
| *Monoamine Oxidase B** | 95 | 51.81 | 539 | 0 |
| *Aromatic Acid Decarboxylase* | 99 | 39.38 | 386 | 4e^-130^ |
| *Aldehyde Dehydrogenase* | 94 | 66.46 | 701 | 0 |

**SI Table 3: Alignment information between *Capitella teleta* and *Caenorhabditis***

***elegans, and Capitella teleta* and *Homo sapiens* for proteins involved in the dopamine pathway.** Alignments were done using NCBI Global Alignments. Percent identity refers to the percentage of the amino acids that are identical between the sequences. Positives refers to the amount of amino acids that are the same or have similar properties between each sequence. Gaps refer to the amount of spaces in the alignment to account for any insertions/deletions in one of the sequences.

| **Species** | **Percent identity** | **Positives (%)** | **Gaps (%)** |
| --- | --- | --- | --- |
| **Dopamine Receptor** | | | |
| *C. teleta* vs. *C. elegans* | 34 | 50 | 16 |
| *C. teleta* vs. *H. sapiens* | 36 | 54 | 7 |
| **Tyrosine Hydroxylase** | | | |
| *C. teleta* vs. *C. elegans* | 72 | 83 | 0 |
| *C. teleta* vs. *H. sapiens* | 56 | 71 | 0 |
| **Dopamine ß-Hydroxylase** | | | |
| *C. teleta* vs. *C. elegans* | 37 | 56 | 6 |
| *C. teleta* vs. *H. sapiens* | 45 | 63 | 1 |
| **Catechol-O-Methyltransferase** | | | |
| *C. teleta* vs. *C. elegans* | - | - | - |
| *C. teleta* vs. *H. sapiens* | 27 | 51 | 2 |
| **Monoamine Oxidase B** | | | |
| *C. teleta* vs. *C. elegans* | 23 | 38 | 12 |
| *C. teleta* vs. *H. sapiens* | 52 | 69 | 0 |
| **Aromatic Acid Decarboxylase** | | | |
| *C. teleta* vs. *C. elegans* | 40 | 58 | 9 |
| *C. teleta* vs. *H. sapiens* | 64 | 79 | 0 |
| **Aldehyde Dehydrogenase** | | | |
| *C. teleta* vs. *C. elegans* | 66 | 83 | 0 |
| *C. teleta* vs. *H. sapiens* | 64 | 80 | 0 |

The differential gene expression analyses revealed that *C. teleta* contain almost all of the transcripts examined in the dopamine pathway. Apart from Cachetol-O-Methyltransferase, the rest of the genes in the pathway were expressed at the juvenile life stage (SI Figure 1). Aldehyde dehydrogenase had the highest expression, followed by aromatic acid decarboxylase, dopamine β-hydroxylase, monoamine oxidase B, and finally, the dopamine receptor, which had the lowest expression (SI Figure 1).

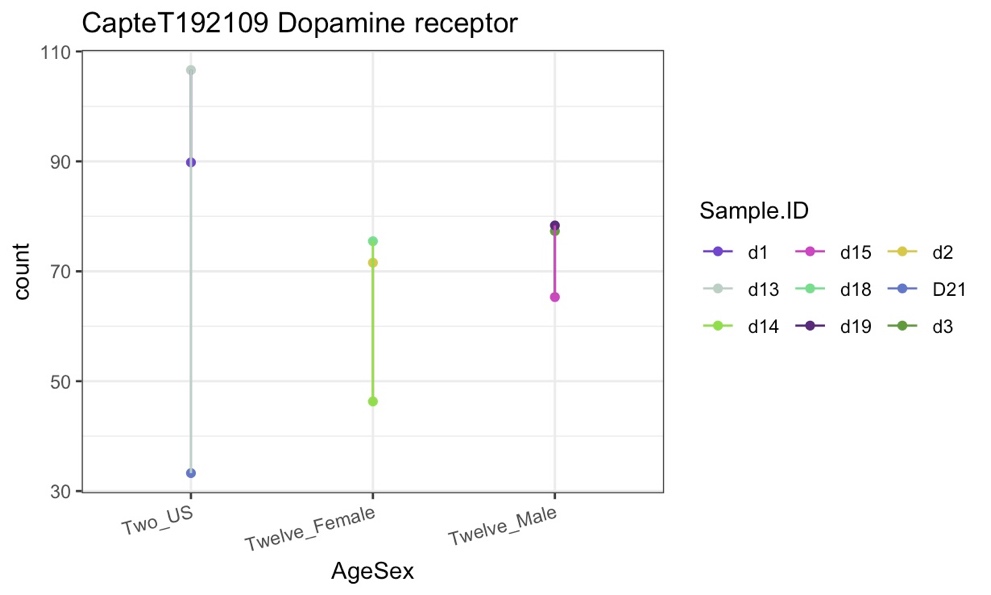

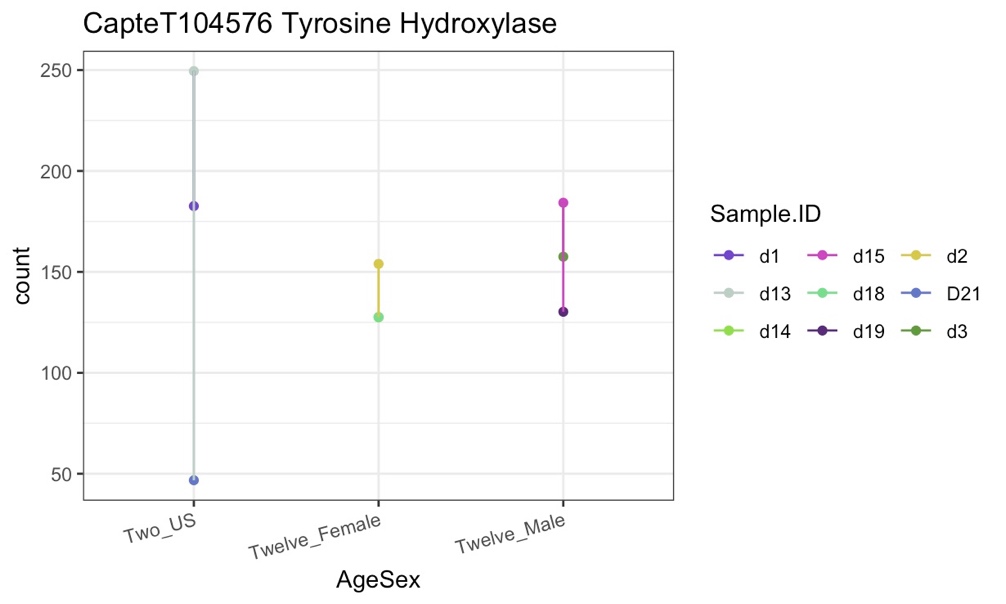

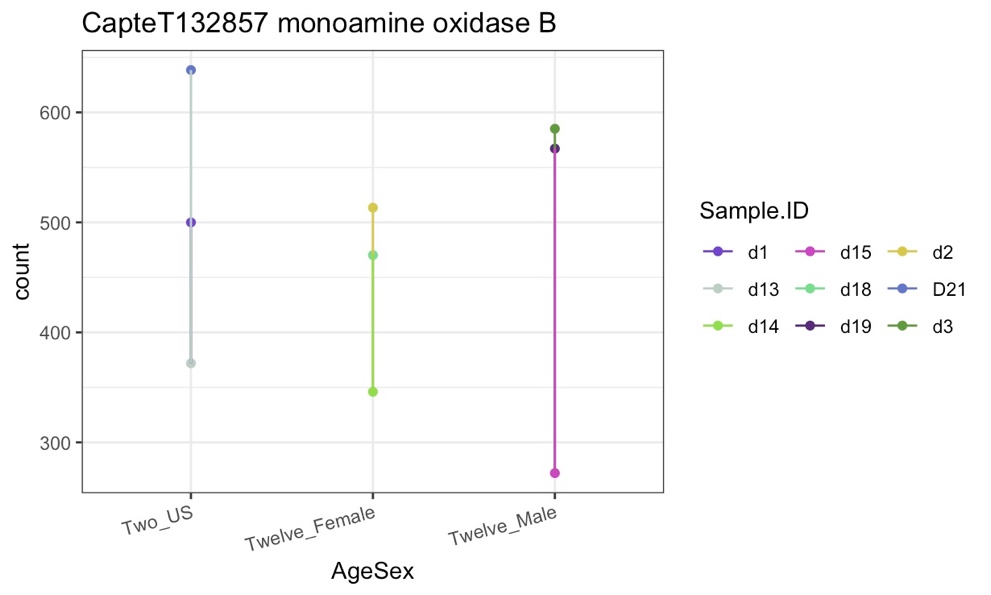

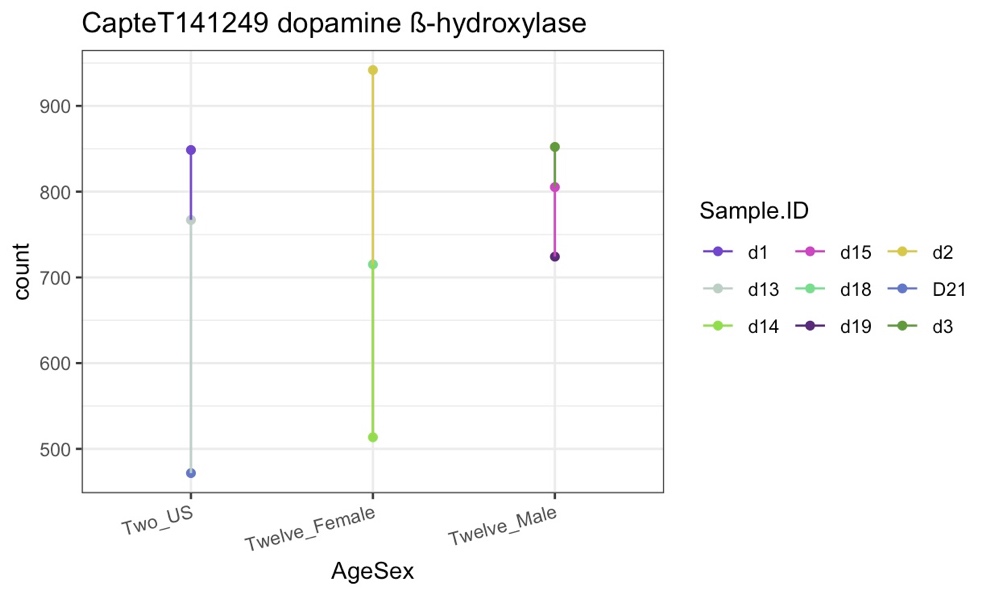

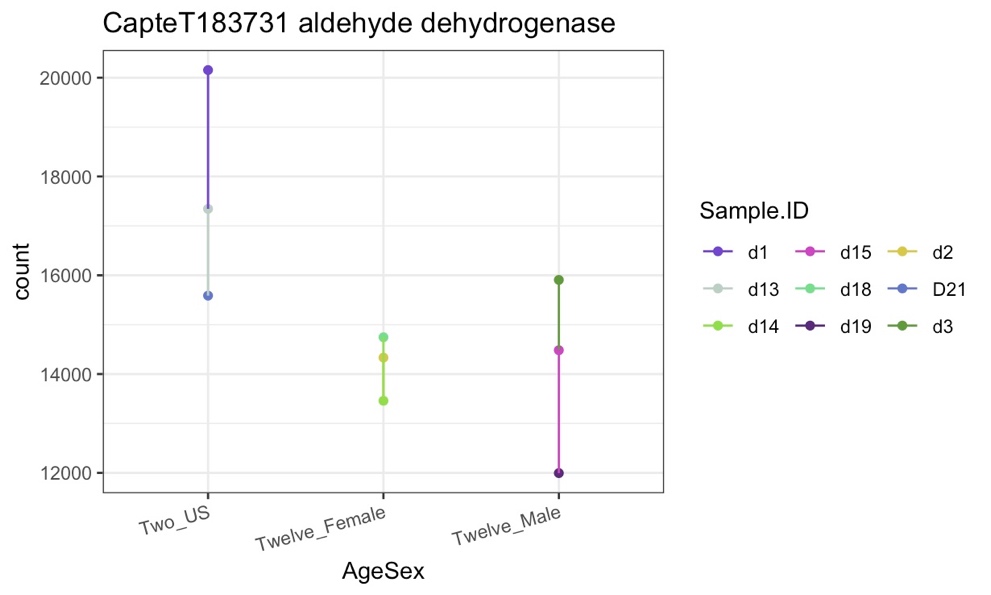

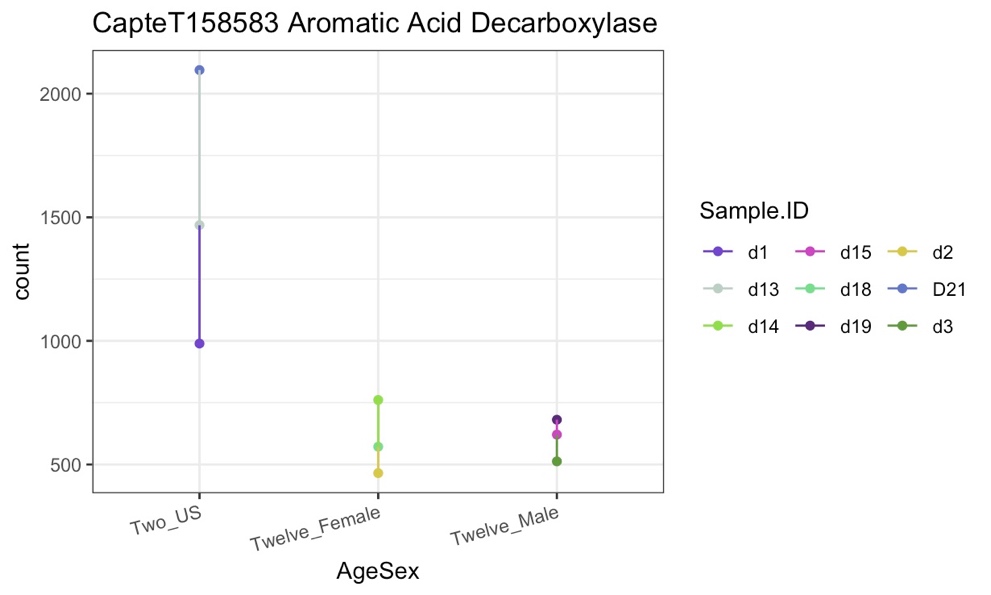

**SI Figure 1: *Expression* counts of genes involved in the dopamine pathway for juveniles, females, and males.** The plots show the dopamine receptor (A), tyrosine hydroxylase (B), monoamine oxidase B (C), dopamine β-hydroxylase (D), aldehyde dehydrogenase (E), and aromatic acid decarboxylase (F). The different groups are Two_US (juveniles), Twelve_Female (females), and Twelve_Male (males). *C.* *teleta* juveniles were sampled at 2 weeks old and the males and females were sampled at 12 weeks old. There were 3 biological replicates for each group, each of a pool of 10 worms and 5 worms for juvenile and adult worms, respectively.

Using InterProScan with the *C. elegans* and *C. teleta,* and *H. sapiens* dopamine receptor sequences, several motifs were found to be conserved between the species, including the “dopamine receptor family”, “rhodopsin-like GPCR superfamily signature”, and G-protein coupled receptors family 1 signature” in very similar locations within the sequences (SI Figure 2, SI Table 4). Additionally, several ligand binding sites were found to be conserved between the sequences as well (SI Figure 2, SI Table 4). Specifically, within the *C. teleta* sequence, additional motifs were found, including the 7tmA_D2-like_dopamine_R domain, G-protein coupled receptors family 1 profile domain, and adrenergic receptor-related G-protein coupled receptor motif (SI Table 4).

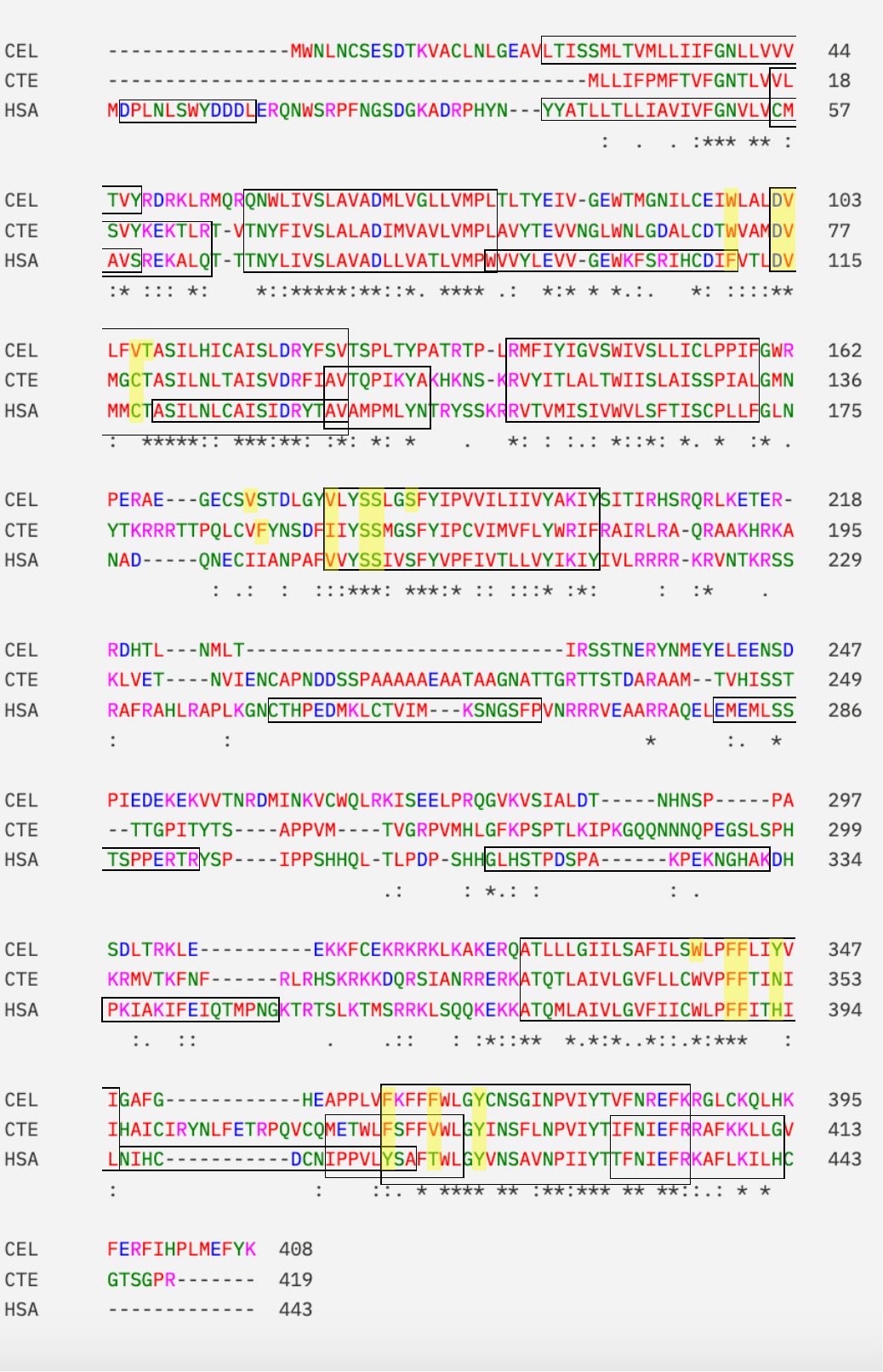

**SI Figure 2: Global alignment of dopamine receptor sequences between *C. teleta, H. sapiens,* and *C. elegans****.* Boxes indicate conserved areas between different sequences. The yellow highlights indicate ligand binding domains. * indicates positions that have a fully conserved residue. : indicates strong conservation between groups that have similar properties. . indicates weak conservation between groups that have similar properties.

**SI Table 4: Motif information for the *Capitella teleta* dopamine receptor.** This information was found using InterProScan. Domains represent functional, structural, or sequence units. Family refers to groups of proteins that share common evolutionary origins, seen as similar functions, sequences, or protein structures. A homologous superfamily represents groups of proteins that share common evolutionary origins, reflected as similarity in the protein structure. Unintegrated represent motifs whose signatures are not classified. Sites refers to short sequences that have one or more conserved residues.

|  | **Amino Acid Numbers** | | | | | | | | | | |
| --- | --- | --- | --- | --- | --- | --- | --- | --- | --- | --- | --- |
| **Motif** | **#1** | **#2** | **#3** | **#4** | **#5** | **#6** | **#7** | **#8** | **#9** | **#10** | **#11** |
| ***Family*** | | | | | | | | | | | |
| G protein-coupled receptor, rhodopsin-like | 6-411 |  |  |  |  |  |  |  |  |  |  |
| Dopamine receptor family | 17-27 | 97-105 | 373-384 | 398-412 |  |  |  |  |  |  |  |
| Rhodopsin-like GPCR superfamily signature | 30-51 | 76-98 | 112-133 | 156-179 | 330-354 | 378-404 |  |  |  |  |  |
| G-protein coupled receptors family 1 signature | 82-98 |  |  |  |  |  |  |  |  |  |  |
| ***Domain*** | | | | | | | | | | | |
| 7tmA_D2-like_dopamine_R | 1-407 |  |  |  |  |  |  |  |  |  |  |
| GPCR, rhodopsin-like, 7TM | 12-396 |  |  |  |  |  |  |  |  |  |  |
| G-protein coupled receptors family 1 profile | 12-396 |  |  |  |  |  |  |  |  |  |  |
| ***Unintegrated*** | | | | | | | | | | | |
| Adrenergic receptor-related G-protein coupled receptor | 1-411 |  |  |  |  |  |  |  |  |  |  |
| Family A G protein-coupled receptor-like | 2-187 | 309-413 |  |  |  |  |  |  |  |  |  |
| ***Sites*** | | | | | | | | | | | |
| Ligand binding site | 72 | 76-77 | 80 | 150 | 156 | 159-160 | 348-349 | 352 | 378 | 382 | 386 |

A phylogenetic tree was made for the dopamine receptor with bootstrap support (SI Figure 3). The species and accession numbers used to construct the tree can be viewed in SI Table 5. Some nodes had lower bootstrap support, making it difficult to determine some placements within the tree. The *C. teleta* sequence clustered close to the *Liolophura japonica* (Pilsbry, 1893), *Octopus sinensis* (D’Orbigny, 1835), and *Mytilus trossulus* (Gould, 1850) sequences (SI Figure 3).

**SI Table 5: Accession numbers for the species and sequences used to make the dopamine receptor phylogenetic tree.** Accession numbers were found using NCBI blastp.

| **Species** | **Accession** |
| --- | --- |
| ***Capitella teleta*** | ELT96301.1 |
| ***Homo sapiens*** | AAC78779.1 |
| ***Caenorhabditis elegans*** | NP_001024569.1 |
| ***Danio rerio*** | NP_001012636.1 |
| ***Xenopus laevis*** | XP_041427336.1 |
| ***Drosophila melanogaster*** | AAN15957.1 |
| ***Lytechinus pictus*** | XP_054766451.2 |
| ***Octopus sinensis*** | XP_036358616.1 |
| ***Liolophura japonica*** | XP_064599039.1 |
| ***Macrobrachium rosenbergii*** | XP_066939829.1 |
| ***Parasteatoda tepidariorum*** | XP_015920481.1 |
| ***Coccinella septempunctata*** | XP_044747683.1 |
| ***Mytilus trossulus*** | XP_063439700.1 |
| ***Anguilla anguilla*** | ABH06893.1 |
| ***Oncorhynchus gorbuscha*** | XP_046178128.1 |
| ***Scyliorhinus canicula*** | XP_038635291.1 |
| ***Caretta caretta*** | XP_048683534.1 |
| ***Alligator sinensis*** | XP_006026722.1 |

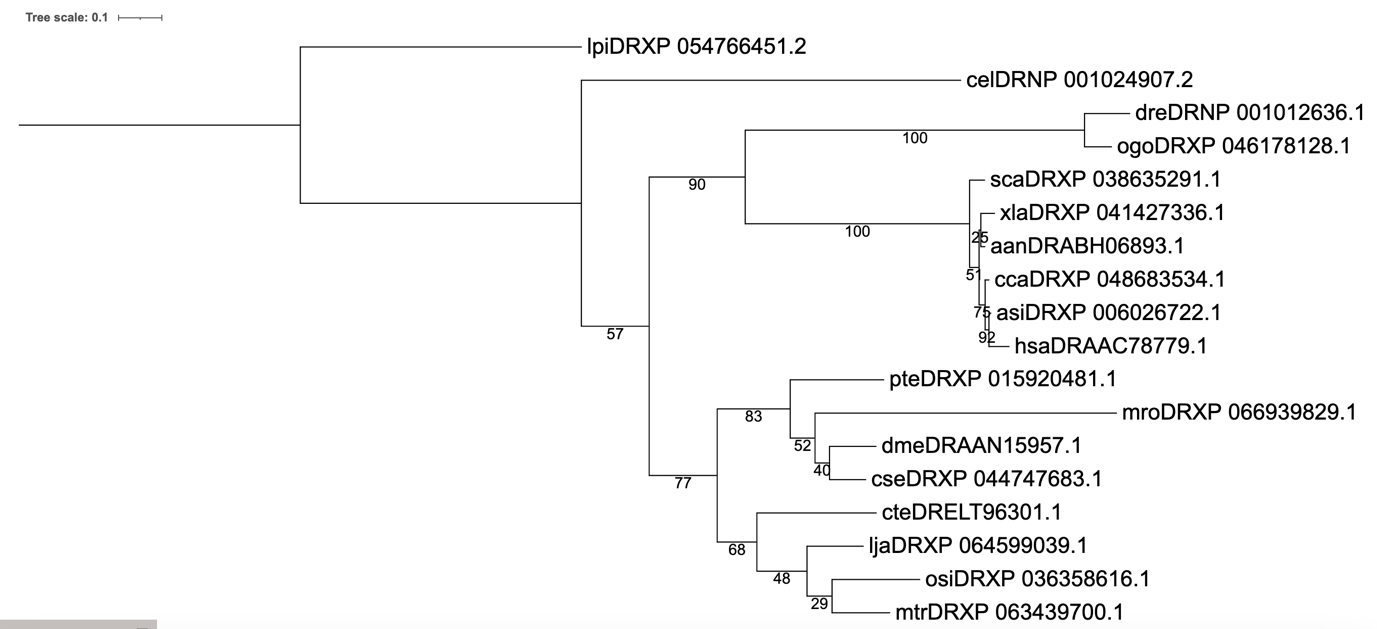

**Figure 3: Phylogenetic tree of dopamine receptor proteins across various species.** Sequences were found using the *C. teleta* dopamine receptor sequence in NCBI blastp, then aligned using Clustal Omega. Alignments were masked using TrimAI and phylogenetic analyses were run using RaxML with 100 bootstraps. A LG+I+G model of amino acid substitution was used for the phylogenetic analyses. Nodes are labelled with the bootstrap support (%; 100 bootstraps). Species acronyms are: aan: *Anguilla anguilla,* Asi: *Alligator sinensis,* cel: *Caenorhabditis elegans,* cca: *Caretta caretta,* cse: *Coccinella septempunctata,* cte: *Capitella teleta,* dme: *Drosophila melanogaster,* dre: *Danio rerio,* has: *Homo sapiens,* lja: *Liolophura japonica,* lpi: *Lytechinus pictus,* mro: *Macrobrachium rosenbergii,* mtr: *Mytilus trossulus,* ogo: *Oncorhynchus gorbuscha*, osi: *Octopus sinensis,* pte: *Parasteatoda tepidariorum,* sca: *Scyliorhinus canicular,* xle: *Xenopus laevis.*

Serotonin

Using the *C. elegans* and *H. sapiens* serotonin transporter, serotonin receptor, tryptophan hydroxylase, aromatic acid decarboxylase, monoamine oxidase A, aralkylamine N-acetyltransferase, acetylserotonin O-methyltransferase, and melatonin receptor sequences, homologues were found in *C. teleta* (accession numbers can be seen in SI Table 6). The *C. teleta* serotonin transporter, serotonin receptor, tryptophan hydroxylase, aromatic acid decarboxylase homologues were found using the *C. elegans* sequences, and monoamine oxidase A, aralkylamine N-acetyltransferase, acetylserotonin O-methyltransferase, and melatonin receptor homologues were found using *H. sapiens* sequences. With the exception of , aralkylamine N-acetyltransferase and acetylserotonin O-methyltransferase, all *C. teleta* homologues were able to align with the *C. elegans* and *H. sapiens* sequences (SI Table 7). The specific values for these BLASTs can be found in SI Table 6. Except for aralkylamine N-acetyltransferase, all BLASTs yielded high max scores, and all had relatively low e-values, high query covers, with percent identities ranging from 30.23-51.82 (SI Table 7). Again, the homologues seemed to align better with the *H. sapiens* sequences, having more positives, fewer gaps, and a higher percent identity (SI Table 8).

**SI Table 6: Accessions for proteins involved in the serotonin pathway from *Capitella teleta, Caenorhabditis elegans,* and *Homo sapiens****.* Accessions were taken off of NCBI.

| **Protein** | **Capitella teleta** | **Caenorhabditis elegans** | **Homo sapiens** |
| --- | --- | --- | --- |
| SERT | ELU16974.1 | NP_491095.3 | AAB26687.1 |
| Serotonin Receptor | ELU07848.1 | NP_497452.1 | BAA94488.1 |
| Tryptophan Hydroxylase | ELU12014.1 | AAD30115.1 | NP_004170.1 |
| Aromatic Acid Decarboxylase | ELU12210.1 | CCD63121.1 | CAG33005.1 |
| Monoamine Oxidase A | ELT97776.1 | NP_001369790.1 | P21397.1 |
| Aralkylamine N-acetyltransferase | ELU18274.1 |  | Q16613.1 |
| Acetylserotonin O-methyltransferase | ELT89989.1 |  | P46597.1 |
| Melatonin Receptor | ELT93417.1 | NP_509725.2 | P48039.1 |

**SI Table 7: BLAST outputs from *Capitella teleta* for proteins involved in the serotonin pathway.** BLASTS were run against *Caenorhabditis elegans* through NCBI blastp. * indicates the BLAST was run between *Capitella teleta* and *Homo sapiens.* Query Cover provides an indication of the length of each subject refers to the length of the target sequence compared to the reference sequence. Percent identity refers to the percentage of the amino acids that are identical between the sequences. Max score refers to the highest score calculated for the alignment between the sequences. The E-value refers to the number of alignments expected by chance with the calculated score.

| **Sequence** | **Query Cover (%)** | **Percent Identity** | **Max Score** | **E-value** |
| --- | --- | --- | --- | --- |
| *SERT* | 80 | 51.82 | 595 | 0 |
| Serotonin Receptor | 85 | 43.26 | 291 | 4e^-95^ |
| Tryptophan Hydroxylase | 78 | 50.46 | 435 | 3e^-149^ |
| Aromatic Acid Decarboxylase | 99 | 39.38 | 386 | 4e^-130^ |
| Monoamine Oxidase A* | 94 | 51.61 | 541 | 0 |
| Aralkylamine N-acetyltransferase* | 71 | 31.33 | 63.2 | 1e^-12^ |
| Acetylserotonin O-methyltransferase* | 97 | 30.23 | 149 | 7e^-42^ |
| Melatonin Receptor* | 82 | 29.35 | 135 | 1e^-36^ |

**SI Table 8: Alignment information between *Capitella teleta* and *Caenorhabditis***

***elegans, and Capitella teleta* and *Homo sapiens* for proteins involved in the serotonin pathway.** Alignments were done using NCBI Global Alignments. Percent identity refers to the percentage of the amino acids that are identical between the sequences. Positives refers to the amount of amino acids that are the same or have similar properties between each sequence. Gaps refer to the amount of spaces in the alignment to account for any insertions/deletions in one of the sequences.

| **Species** | **Percent identity** | **Positives (%)** | **Gaps (%)** |
| --- | --- | --- | --- |
| **SERT** | | | |
| *C. teleta* vs. *C. elegans* | 52 | 72 | 2 |
| *C. teleta* vs. *H. sapiens* | 56 | 75 | 2 |
| **Serotonin Receptor** | | | |
| *C. teleta* vs. *C. elegans* | 42 | 68 | 10 |
| *C. teleta* vs. *H. sapiens* | 43 | 61 | 4 |
| **Tryptophan Hydroxylase** | | | |
| *C. teleta* vs. *C. elegans* | 50 | 66 | 8 |
| *C. teleta* vs. *H. sapiens* | 57 | 74 | 5 |
| **Aromatic Acid Decarboxylase** | | | |
| *C. teleta* vs. *C. elegans* | 40 | 58 | 9 |
| *C. teleta* vs. *H. sapiens* | 64 | 79 | 0 |
| **Monoamine Oxidase A** | | | |
| *C. teleta* vs. *C. elegans* | 23 | 38 | 12 |
| *C. teleta* vs. *H. sapiens* | 52 | 70 | 0 |
| **Aralkylamine N-acetyltransferase** | | | |
| *C. teleta* vs. *C. elegans* | - | - | - |
| *C. teleta* vs. *H. sapiens* | 31 | 45 | 2 |
| **Acetylserotonin O-methyltransferase** | | | |
| *C. teleta* vs. *C. elegans* | - | - | - |
| *C. teleta* vs. *H. sapiens* | 30 | 50 | 9 |
| **Melatonin Receptor** | | | |
| *C. teleta* vs. *C. elegans* | 26 | 46 | 15 |
| *C. teleta* vs. *H. sapiens* | 29 | 48 | 10 |

The differential gene expression analyses showed that *C. teleta* contain almost all of the proteins examined in the serotonin pathway. All the transcripts looked juvenile life stage for the serotonin pathway were present, except for the melatonin receptor, and acetylserotonin o-methyltransferase was very lowly expressed in the juvenile life stage, but above the cutoff point of 10 counts (SI Figure 4). Aromatic acid decarboxylase had the highest expression, followed by the SERT, monoamine oxidase A, aralkylamine n-acetyltransferase, serotonin receptor, tryptophan hydroxylase, and then acetylserotonin o-methyltransferase (SI Figure 4)

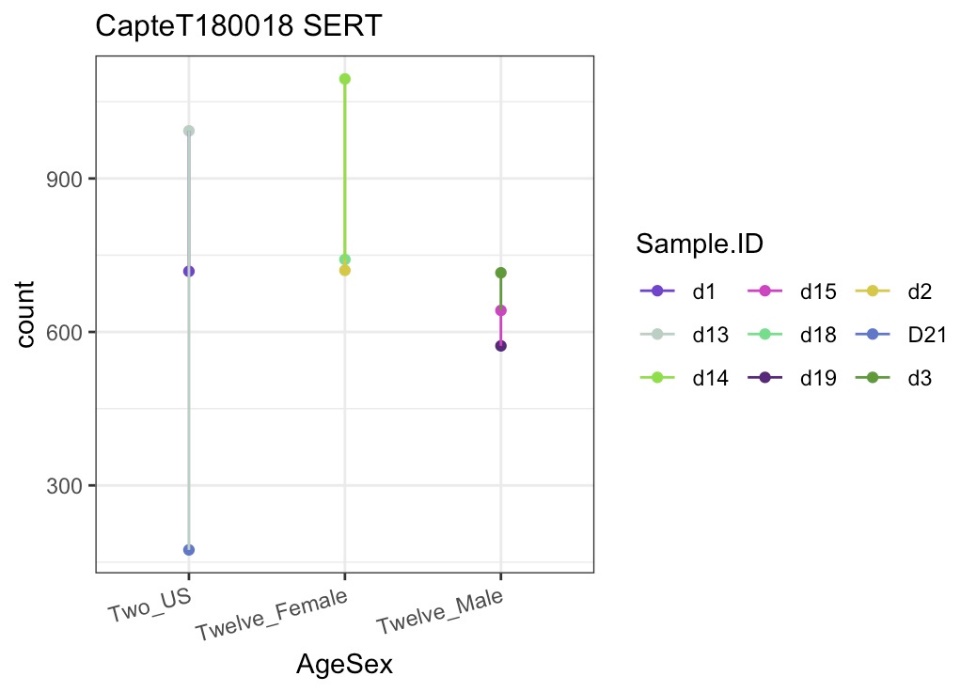

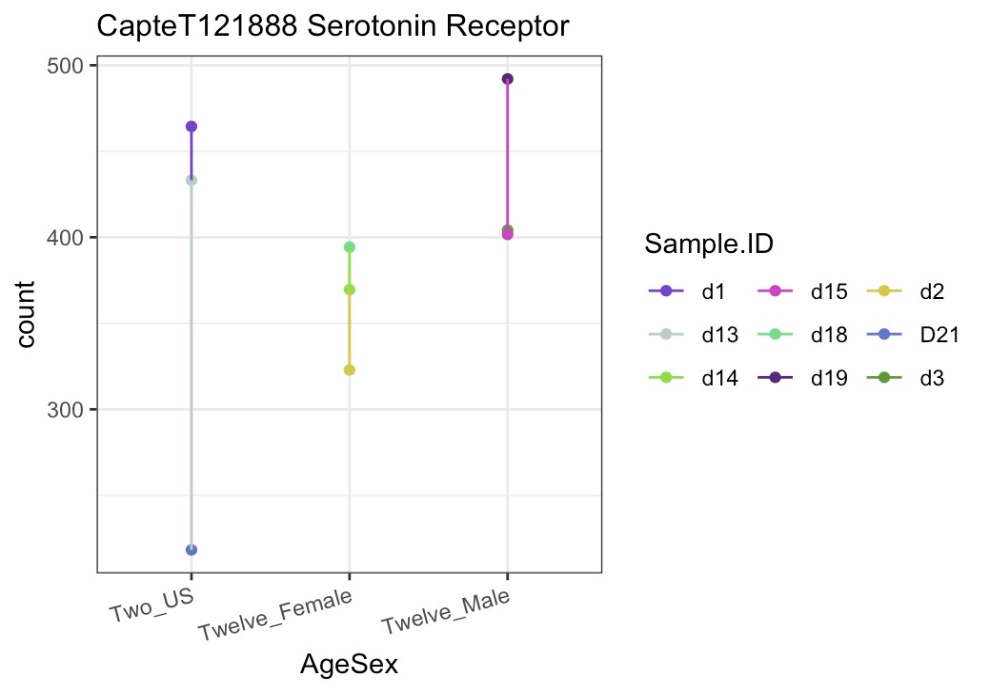

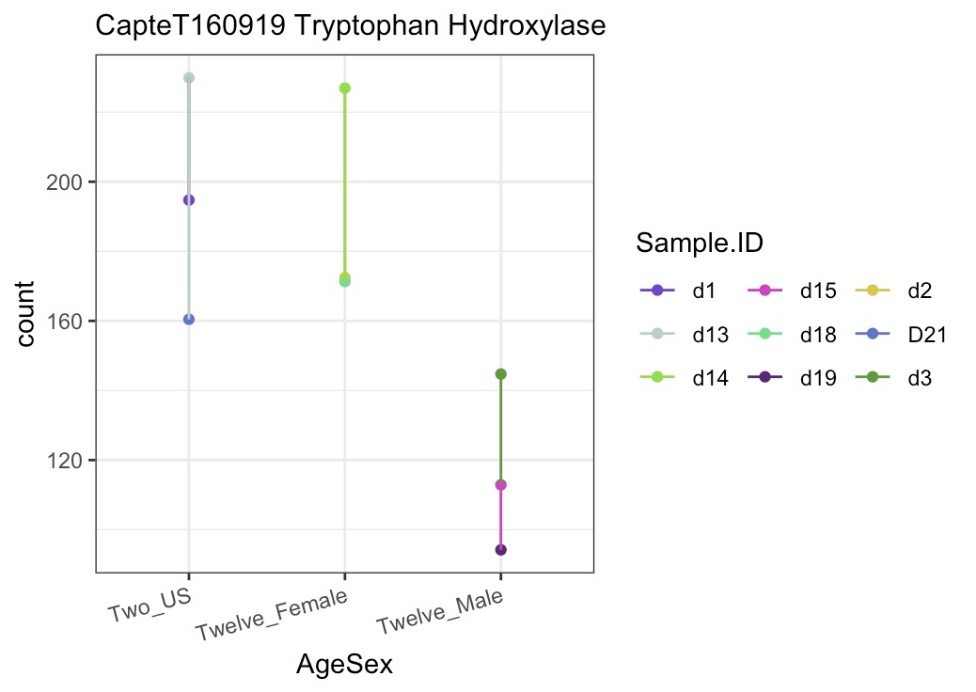

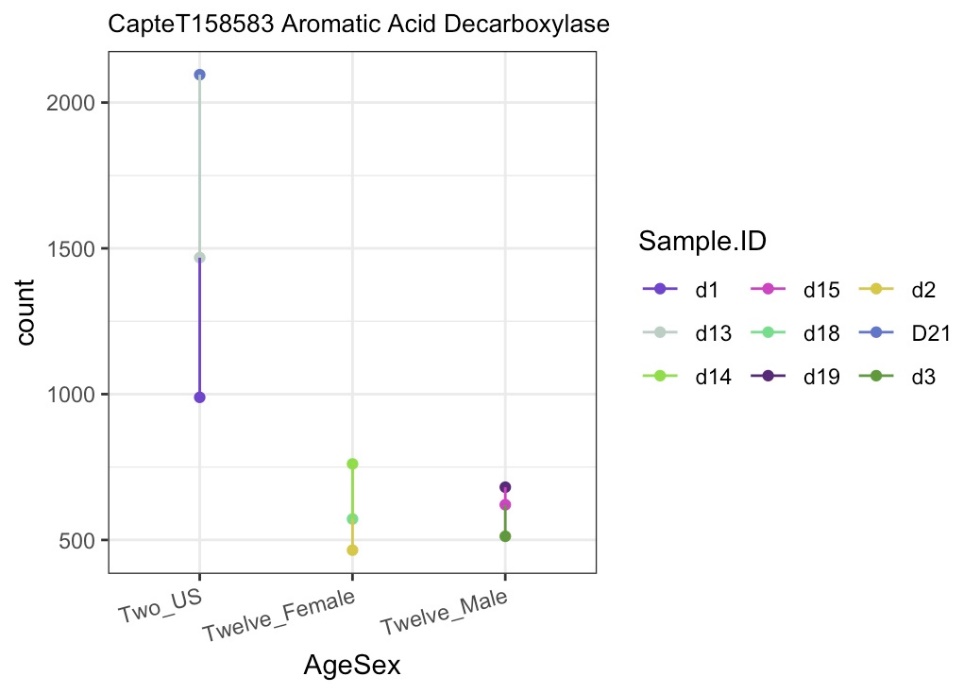

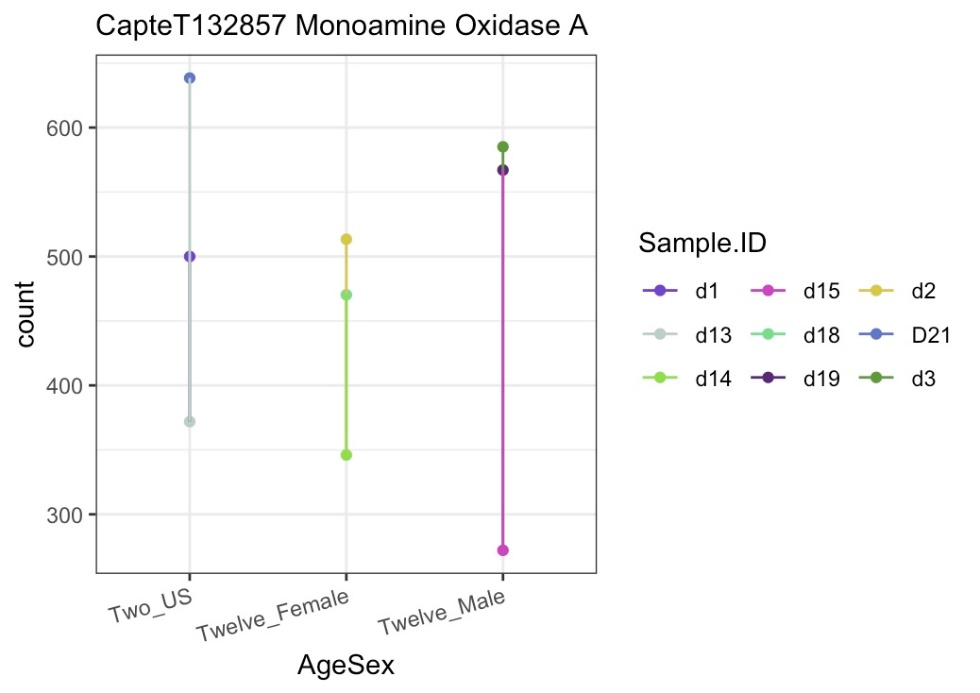

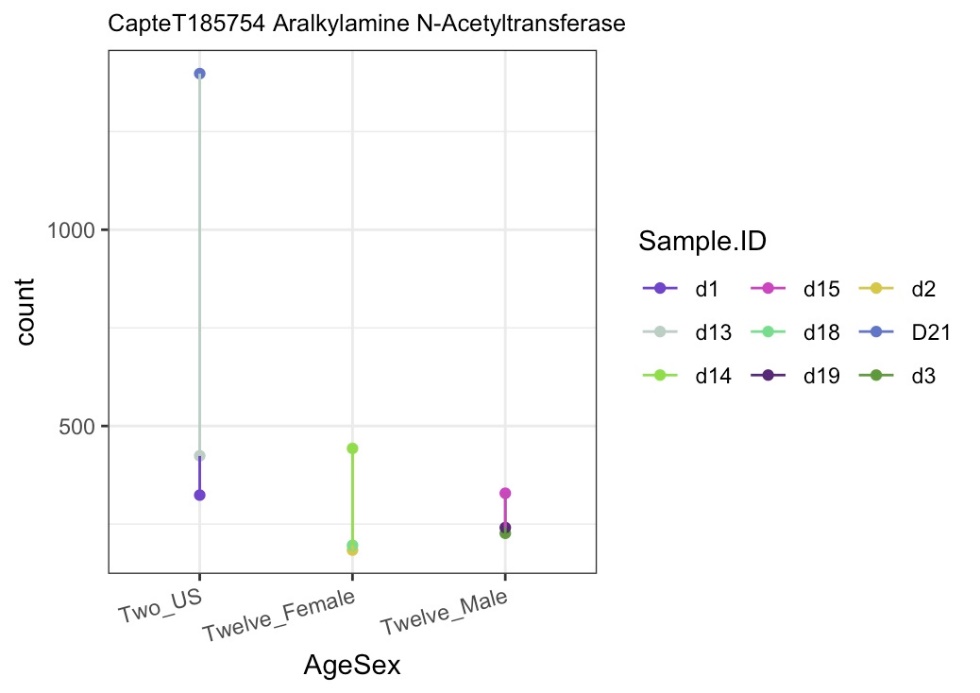

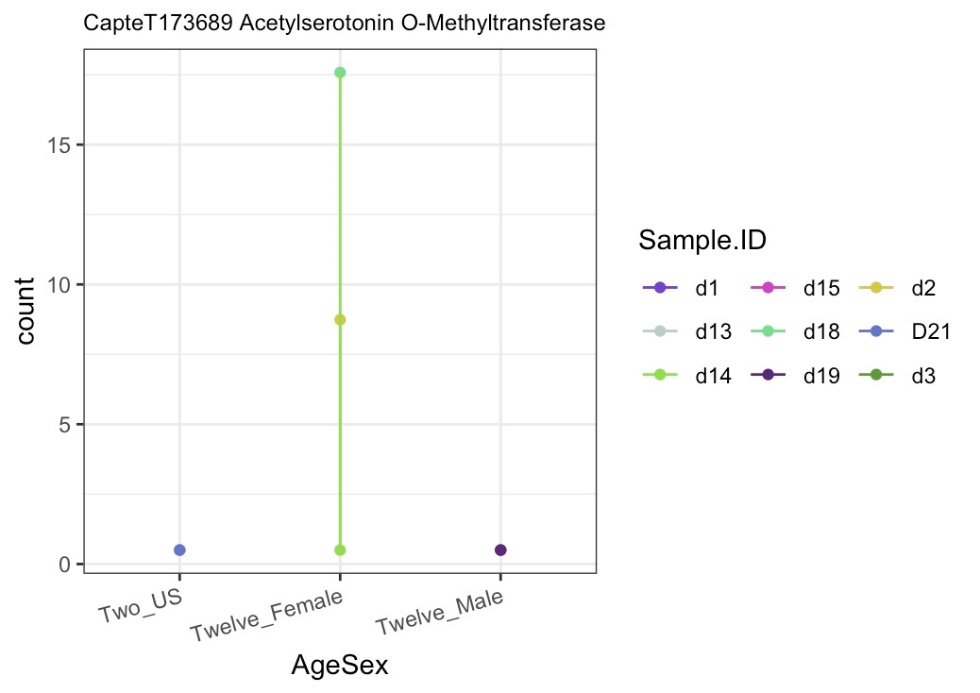

**SI Figure 4: Expression counts of proteins involved in the serotonin pathway for juvenile, female, and male *Capitella teleta*.** The plots show the SERT (A), serotonin receptor (B), tryptophan hydryoxylase (C), aromatic acid decarboxylase (D), monoamine oxidase A (E), aralkylamine n-acetyltransferase (F), and acetylserotonin o-methyltransferase (G). The different groups are Two_US (juveniles), Twelve_Female (females), and Twelve_Male (males). *C.* *teleta* juveniles were sampled at 2 weeks old and the males and females were sampled at 12 weeks old. There were 3 biological replicates for each group, each of a pool of 10 worms and 5 worms for juvenile and adult worms, respectively.

InterProScan was also used with the *C. elegans* and *C. teleta,* and *H. sapiens* SERT sequences. There were again many similar motifs in similar locations throughout the sequence, such as the “sodium:neurotransmitter symporter”, “sodium/chloride neurotransmitter symporter signature” in all 3 species. Ligand binding sites,and Na^+^ binding sites were also found to be consistant across all 3 sequences (SI Figure 5, SI Table 9). Within the *C. teleta* sequence, motifs such as the “SLC6sbd_SERT-like” domain, “sodium:neurotransmitter symporter superfamily”, and “sodium-dependent transporter” were found (SI Table 9).

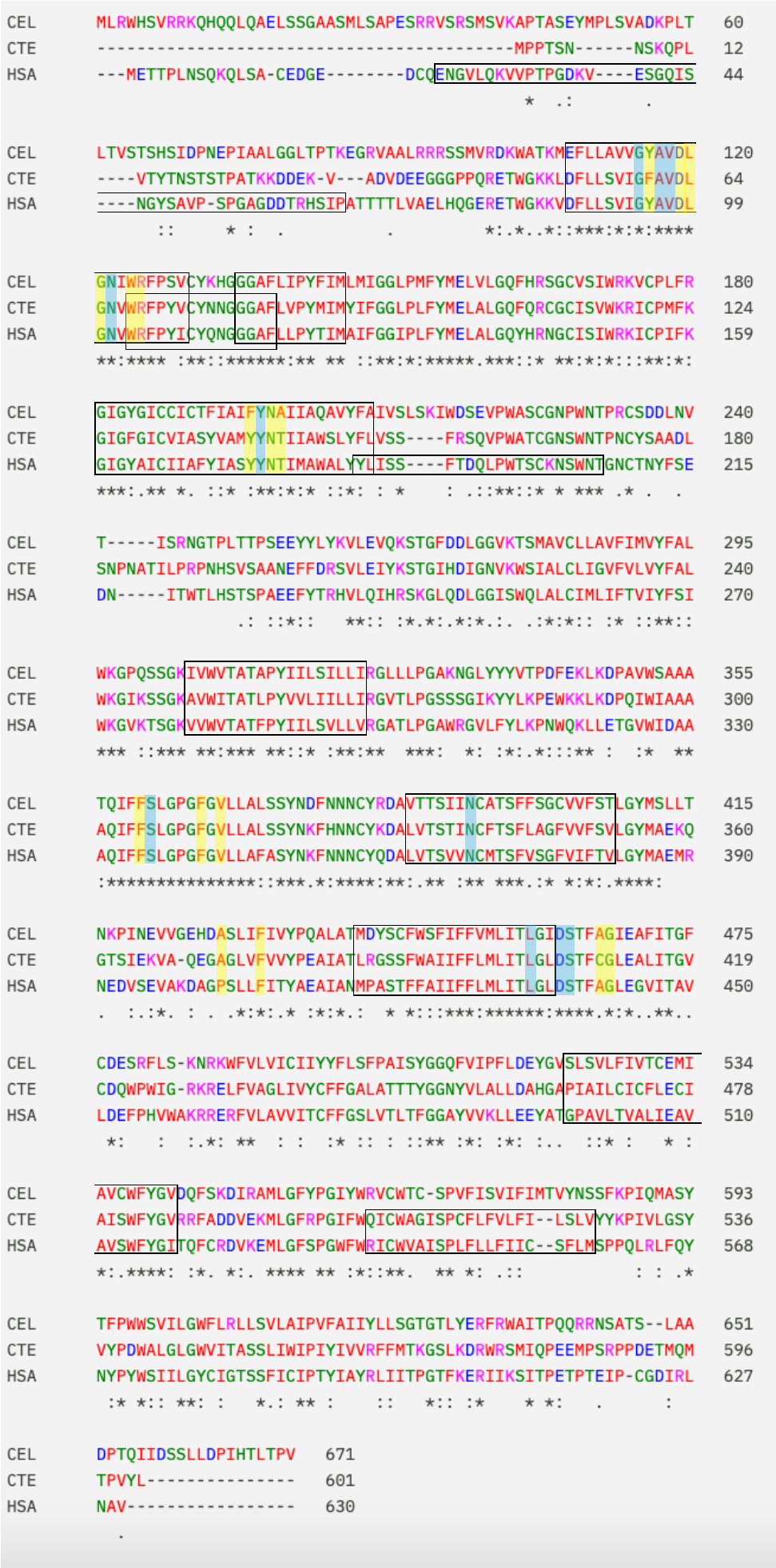

**SI Figure 5: Global alignment of SERT sequences between *C. teleta, H. sapiens,* and *C. elegans****.* Boxes indicate conserved areas between different sequences. The yellow highlights indicate ligand binding domains. The blue highlights indicate sodium binding domains. * indicates positions that have a fully conserved residue. : indicates strong conservation between groups that have similar properties. . indicates weak conservation between groups that have similar properties.

**SI Table 9: Motif information for the *Capitella teleta* SERT.** This information was found using InterProScan. Domains represent functional, structural, or sequence units. Family refers to groups of proteins that share common evolutionary origins, seen as similar functions, sequences, or protein structures. A homologous superfamily represents groups of proteins that share common evolutionary origins, reflected as similarity in the protein structure. Unintegrated represent motifs whose signatures are not classified. Sites refers to short sequences that have one or more conserved residues.

|  | **Amino Acid Numbers** | | | | | | | | | | | | |
| --- | --- | --- | --- | --- | --- | --- | --- | --- | --- | --- | --- | --- | --- |
| **Motif** | **#1** | **#2** | **#3** | **#4** | **#5** | **#6** | **#7** | **#8** | **#9** | **#10** | **#11** | **#12** | **#13** |
| ***Domains*** | | | | | | | | | | | | | |
| SLC6sbd_SERT-like | 44-583 |  |  |  |  |  |  |  |  |  |  |  |  |
| ***Family*** | | | | | | | | | | | | | |
| Sodium:neurotransmitter symporter | 27-584 |  |  |  |  |  |  |  |  |  |  |  |  |
| Sodium:neurotransmitter symporter family signature 1 | 68-82 |  |  |  |  |  |  |  |  |  |  |  |  |
| Sodium/chloride dependent transporter | 27-584 |  |  |  |  |  |  |  |  |  |  |  |  |
| Sodium:neurotransmitter symporter family | 44-566 |  |  |  |  |  |  |  |  |  |  |  |  |
| Sodium:neurotransmitter symporter family profile | 43-570 |  |  |  |  |  |  |  |  |  |  |  |  |
| Sodium/chloride neurotransmitter symporter signature | 52-73 | 81-100 | 125-151 | 250-267 | 332-352 | 386-405 | 466-486 | 506-526 |  |  |  |  |  |
| ***Homologous Superfamily*** | | | | | | | | | | | | | |
| Sodium:neurotransmitter symporter superfamily | 44-565 |  |  |  |  |  |  |  |  |  |  |  |  |
| ***Unintegrated*** | | | | | | | | | | | | | |
| sodium-dependent transporter | 51-441 |  |  |  |  |  |  |  |  |  |  |  |  |
| ***Sites*** | | | | | | | | | | | | | |
| Ligand binding domain | 60-61 | 63-66 | 68-69 | 140-141 | 144 | 147 | 305-306 | 311 | 313 | 372 | 376 | 407 | 410-411 |
| Na binding domain | 59 | 61-62 | 66 | 306 | 338 | 403 | 406-407 |  |  |  |  |  |  |

A phylogenetic tree was made for the SERT with bootstrap support (SI Figure 6). The species and accession numbers used to construct the tree can be viewed in SI Table 10. Some nodes had lower bootstrap support, making it difficult to determine some placements within the tree. The *C. teleta* sequence clustered close to the *L. japonica, O. sinensis*, and *M. trossulus* sequences (SI Figure 6).

**SI Table 10: Accession numbers for the species and sequences used to make the dopamine receptor phylogenetic tree.** Accession numbers were found using NCBI blastp.

| **SERT** | **Accession** |
| --- | --- |
| ***Capitella teleta*** | ELU16974.1 |
| ***Homo sapiens*** | AAB26687.1 |
| ***Caenorhabditis elegans*** | NP_491095.3 |
| ***Danio rerio*** | NP_001170930.1 |
| ***Xenopus Laevis*** | XP_018079948.1 |
| ***Drosophila melanogaster*** | NP_001369117.1 |
| ***Lytechinus pictus*** | XP_054766756.2 |
| ***Ciona intestinalis*** | XP_002125543.1 |
| ***Octopus sinensis*** | XP_036359789.1 |
| ***Liolophura japonica*** | XP_064612763 |
| ***Macrobrachium rosenbergii*** | XP_066975304 |
| ***Parasteatoda tepidariorum*** | XP_042899384.1 |
| ***Coccinella septempunctata*** | XP_044749690.1 |
| ***Mytilus trossulus*** | XP_063396380 |
| ***Anguilla anguilla*** | XP_035292066.1 |
| ***Oncorhynchus gorbuscha*** | XP_046204527 |
| ***Scyliorhinus canicula*** | XP_038673873 |
| ***Caretta caretta*** | XP_048677905.1 |
| ***Alligator sinensis*** | XP_006015335.1 |

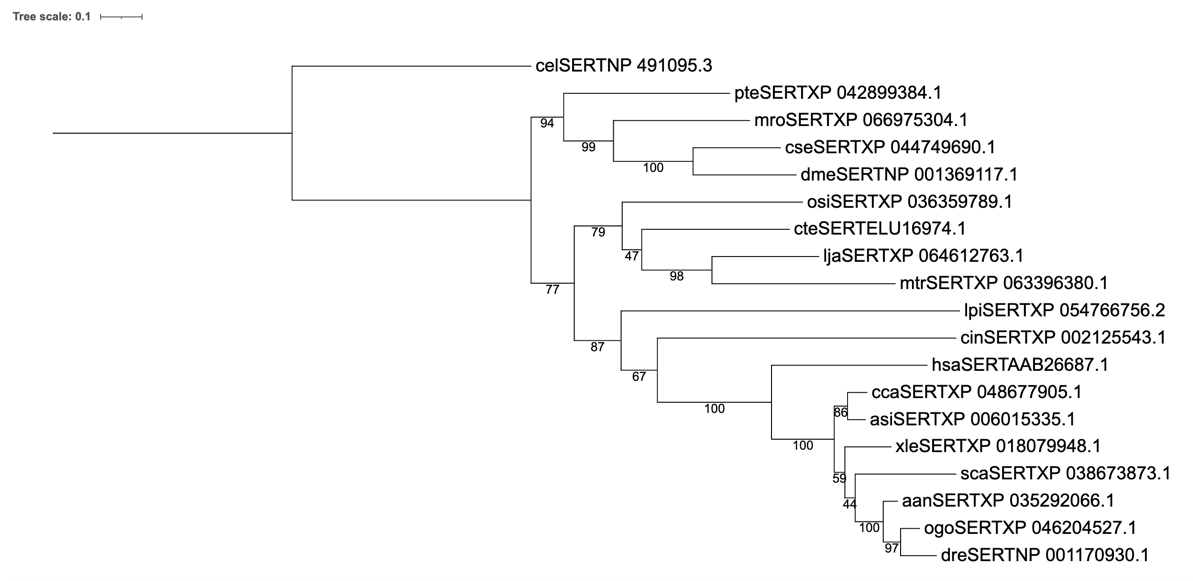

**Figure 6: Phylogenetic tree of SERT proteins across various species.** Sequences were found using the *C. teleta* SERT sequence in NCBI blastp, then aligned using Clustal Omega. Alignments were masked using TrimAI and phylogenetic analyses were run using RaxML with 100 bootstraps. A LG4X+I model of amino acid substitution was used for the phylogenetic analyses. Nodes are labelled with the bootstrap support (%; 100 bootstraps). Species acronyms are: aan: *Anguilla anguilla,* Asi: *Alligator sinensis,* cel: *Caenorhabditis elegans,* cca: *Caretta caretta,* cin: *Ciona intestinalis*, cse: *Coccinella septempunctata,* cte: *Capitella teleta,* dme: *Drosophila melanogaster,* dre: *Danio rerio,* has: *Homo sapiens,* lja: *Liolophura japonica,* lpi: *Lytechinus pictus,* mro: *Macrobrachium rosenbergii,* mtr: *Mytilus trossulus,* ogo: *Oncorhynchus gorbuscha*, osi: *Octopus sinensis,* pte: *Parasteatoda tepidariorum,* sca: *Scyliorhinus canicular,* xle: *Xenopus laevis.*

Acetylcholine

Using the *C. elegans* and *H. sapiens* acetylcholine transporter, acetylcholinesterase, choline acetyltransferase, ATP citrate lyase, and acyl-CoA synthetase, homologues were found in *C. teleta* (accession numbers can be seen in SI Table 11). The specific values for these BLASTs can be found in SI Table 12. All *C. teleta* homologues were found using the *C. elegans* sequences, except for choline acetyltransferase, which was found using the *H. sapiens* sequence. All BLASTs had low e-values, with choline acetyltransferase, ATP citrate lyase, and acyl-CoA synthetase having e-values of 0 (SI Table 12). The acetylcholine receptor had the lowest query cover of 56%, but all other homologues were ~80-100 (SI Table 12). The acetylcholine receptor also had the lowest max score of 238, whereas ATP citrate lyase had the highest at 1368 (SI Table 12). All BLASTs had a moderate percent identity around 50 (SI Table 12). All *C. teleta* homologues were able to align with the *C. elegans* and *H. sapiens* sequences (SI Table 13). The homologues mostly seemed to align better with the *H. sapiens* sequences, having more positives, fewer gaps, and a higher percent identity, except for the acetylcholine receptor, which aligned better with the *C. elegans* sequence (SI Table 13).

**SI Table 11: Accessions for proteins involved in the acetylcholine pathway from *Capitella teleta, Caenorhabditis elegans,* and *Homo sapiens****.* Accessions were taken off of NCBI.

| **Protein** | **Capitella teleta** | **Caenorhabditis elegans** | **Homo sapiens** |
| --- | --- | --- | --- |
| Acetylcholine Receptor | ELT95769.1 | NP_001024236.1 | P11229.2 |
| Acetylcholinesterase | ELU02276.1 | NP_510660.1 | P22303.1 |
| Choline Acetyltransferase* | ELU18773.1 | NP_510624.1 | P28329.4 |
| ATP citrate lyase | ELU08772.1 | NP_508280.1 | P53396.3 |
| acyl-CoA synthetase short chain family member 1 | ELU08878.1 | NP_001021206.1 | Q9NUB1.2 |

**SI Table 12: BLAST outputs from *Capitella teleta* for proteins involved in the acetylcholine pathway.** BLASTS were run against *Caenorhabditis elegans* through NCBI blastp. Query Cover provides an indication of the length of each subject refers to the length of the target sequence compared to the reference sequence. Percent identity refers to the percentage of the amino acids that are identical between the sequences. Max score refers to the highest score calculated for the alignment between the sequences. The E-value refers to the number of alignments expected by chance with the calculated score.

| **Sequence** | **Query Cover (%)** | **Percent Identity** | **Max Score** | **E-value** |
| --- | --- | --- | --- | --- |
| Acetylcholine Receptor | 56 | 49.41 | 238 | 7e^-73^ |
| Acetylcholinesterase | 89 | 42.07 | 446 | 1e^-149^ |
| Choline Acetyltransferase | 79 | 45.69 | 548 | 0 |
| ATP citrate lyase | 99 | 61.57 | 1368 | 0 |
| acyl-CoA synthetase short chain family member 1 | 91 | 61.21 | 845 | 0 |

**SI Table 13: Alignment information between *Capitella teleta* and *Caenorhabditis***

***elegans, and Capitella teleta* and *Homo sapiens* for proteins involved in the acetylcholine pathway.** Alignments were done using NCBI Global Alignments. Percent identity refers to the percentage of the amino acids that are identical between the sequences. Positives refers to the amount of amino acids that are the same or have similar properties between each sequence. Gaps refer to the amount of spaces in the alignment to account for any insertions/deletions in one of the sequences.

| **Species** | **Percent identity** | **Positives (%)** | **Gaps (%)** |
| --- | --- | --- | --- |
| **Acetylcholine Receptor** | | | |
| *C. teleta* vs. *C. elegans* | 49 | 64 | 9 |
| *C. teleta* vs. *H. sapiens* | 46 | 60 | 5 |
| **Acetylcholinesterase** | | | |
| *C. teleta* vs. *C. elegans* | 42 | 58 | 2 |
| *C. teleta* vs. *H. sapiens* | 43 | 59 | 5 |
| **Choline Acetyltransferase** | | | |
| *C. teleta* vs. *C. elegans* | 32 | 59 | 9 |
| *C. teleta* vs. *H. sapiens* | 46 | 62 | 7 |
| **ATP citrate lyase** | | | |
| *C. teleta* vs. *C. elegans* | 61 | 75 | 4 |
| *C. teleta* vs. *H. sapiens* | 71 | 82 | 1 |
| **acyl-CoA synthetase short chain family member 1** | | | |
| *C. teleta* vs. *C. elegans* | 46 | 62 | 4 |
| *C. teleta* vs. *H. sapiens* | 61 | 76 | 0 |

The differential gene expression analyses showed that *C. teleta* contain all of the proteins examined in the acetylcholine pathway. ATP citrate lyase had the highest expression, followed by acetylcholinesterase, acyl-CoA synthetase, acetylcholine receptor, and then choline acetyltransferase (SI Figure 7).

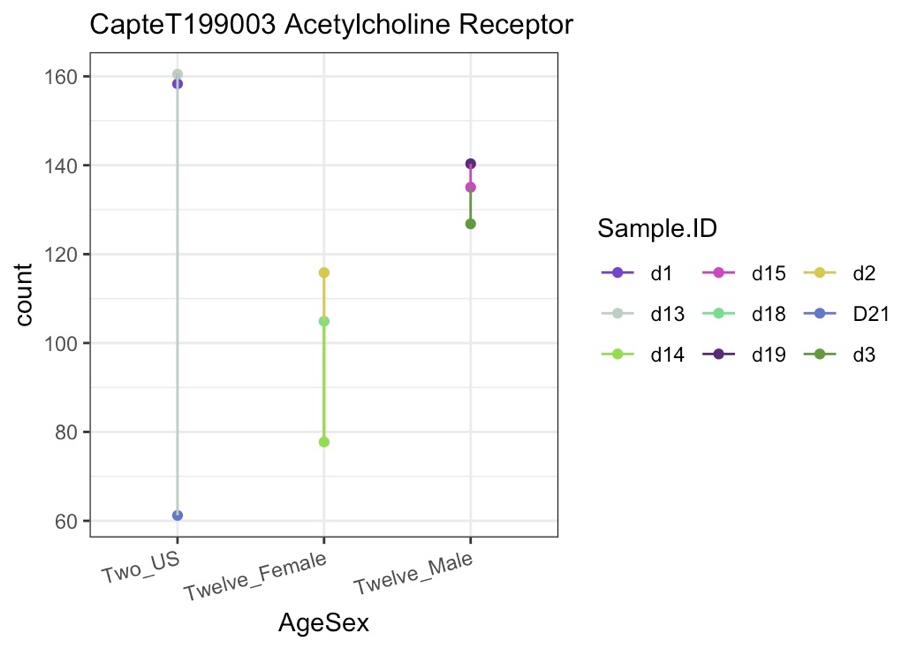

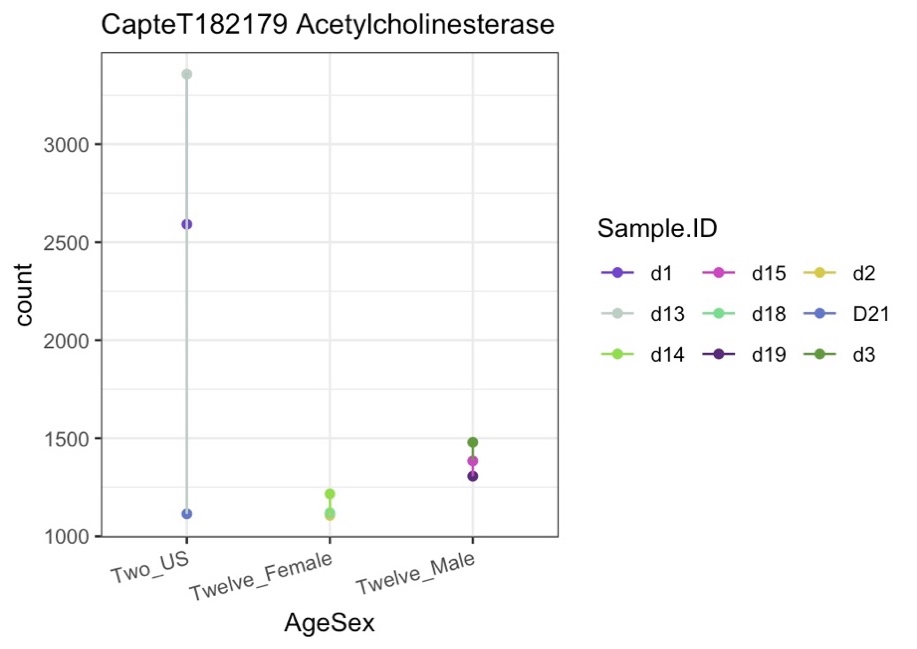

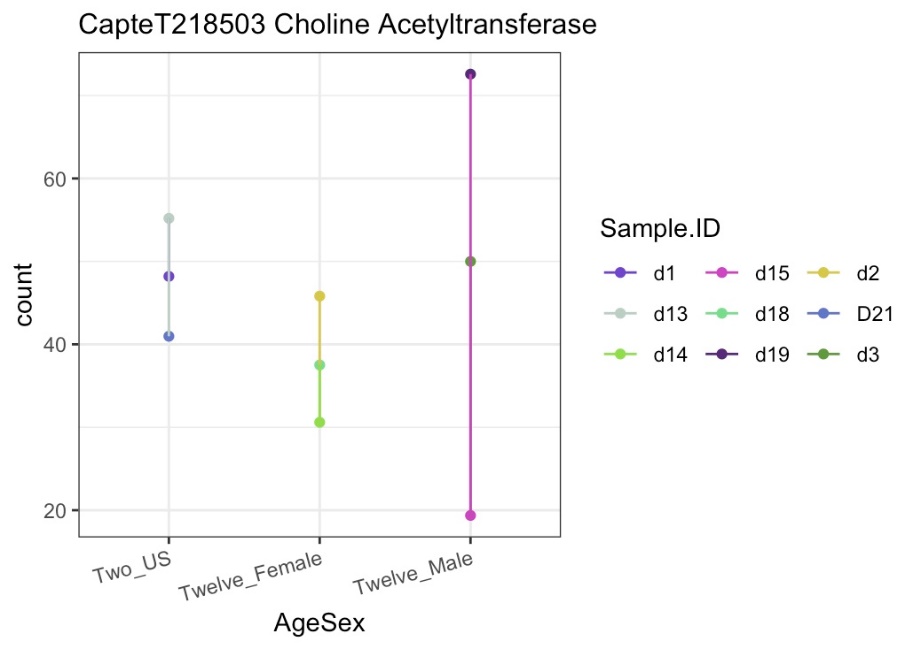

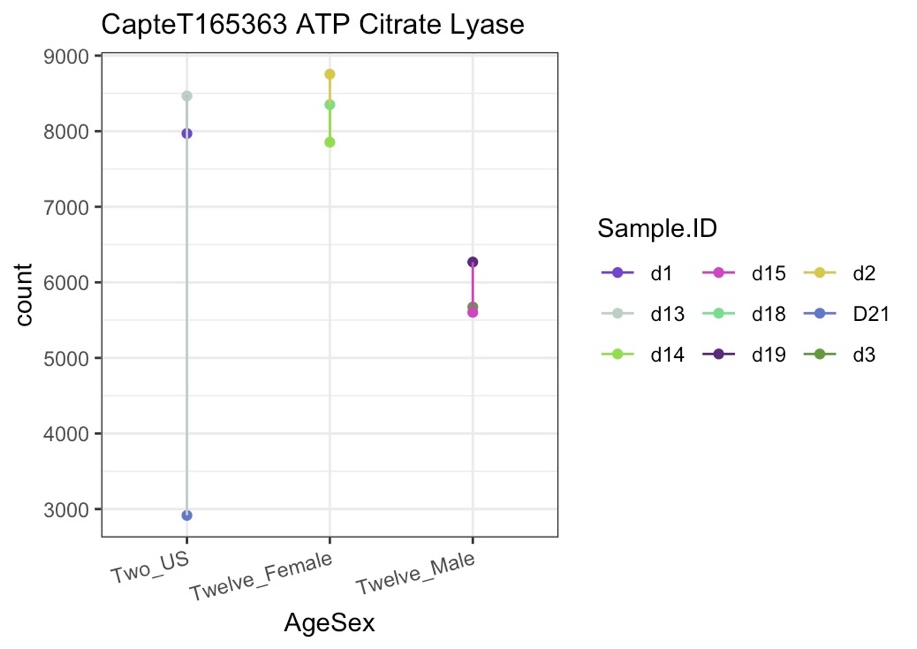

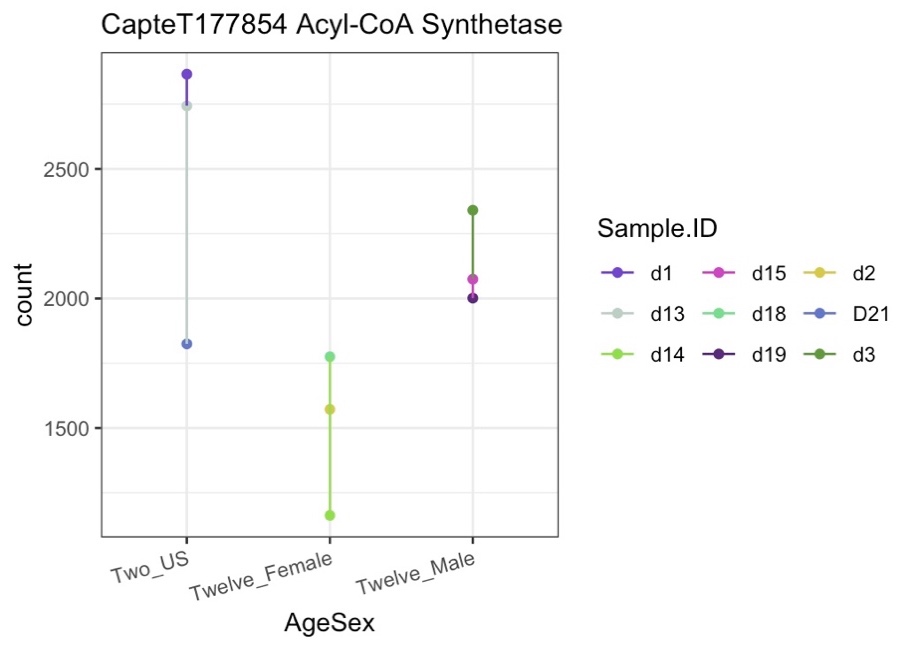

**SI Figure 7: Expression counts of proteins involved in the acetylcholine pathway for juvenile, female, and male *Capitella teleta*.** The plots show the acetylcholine receptor (A), acetylcholinesterase (B), choline acetyltransferase (C), ATP citrate lyase (D), acyl-CoA synthetase (E). The different groups are Two_US (juveniles), Twelve_Female (females), and Twelve_Male (males). *C.* *teleta* juveniles were sampled at 2 weeks old and the males and females were sampled at 12 weeks old. There were 3 biological replicates for each group, each of a pool of 10 worms and 5 worms for juvenile and adult worms, respectively.

InterProScan was also used with the *C. elegans* and *C. teleta,* and *H. sapiens* acetylcholine receptor sequences. There were several similar motifs in similar locations throughout the sequence, such as the “muscarinic acetylcholine receptor”, “GPCR Rhodopsin”, “7tmA_mAChR” domains, and ligand binding sites (SI Figure 8, SI Table 14). Specifically, within the *C. teleta* sequence, family signatures such as the “Rhodopsin-like GPCR superfamily signature” and “G-protein coupled receptors family 1 signature” were found (SI Figure 14).

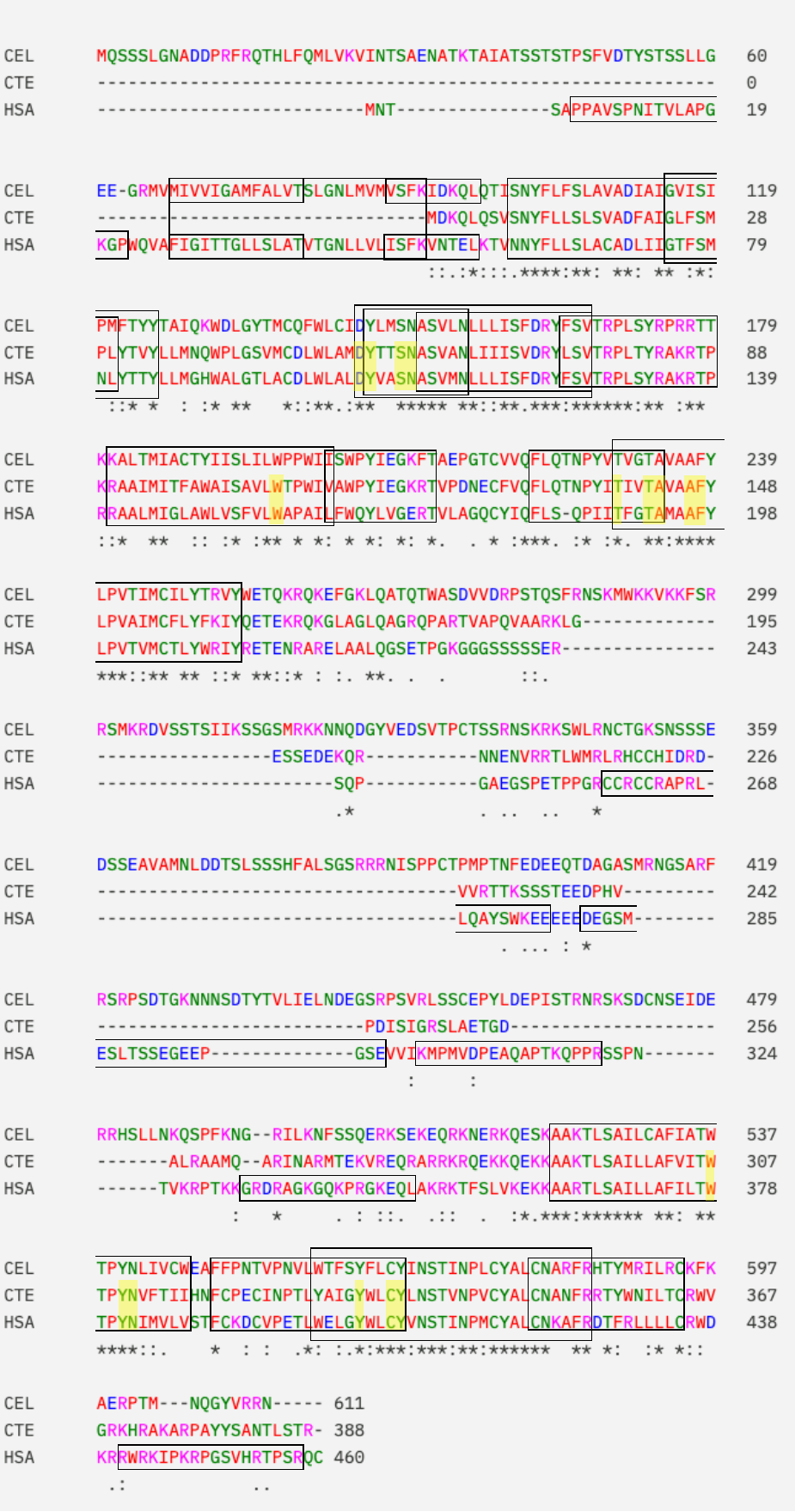

**SI Figure 8: Global alignment of acetylcholine receptor sequences between *C. teleta, H. sapiens,* and *C. elegans****.* Boxes indicate conserved areas between different sequences. The yellow highlights indicate ligand binding domains. * indicates positions that have a fully conserved residue. : indicates strong conservation between groups that have similar properties. . indicates weak conservation between groups that have similar properties.

**SI Table 14: Motif information for the *Capitella teleta* acetylcholine receptor.** This information was found using InterProScan. Domains represent functional, structural, or sequence units. Family refers to groups of proteins that share common evolutionary origins, seen as similar functions, sequences, or protein structures. A homologous superfamily represents groups of proteins that share common evolutionary origins, reflected as similarity in the protein structure. Unintegrated represent motifs whose signatures are not classified. Sites refers to short sequences that have one or more conserved residues.

| **Amino Acid Numbers** | | | | | | | | | | |
| --- | --- | --- | --- | --- | --- | --- | --- | --- | --- | --- |
| **Motif** | **#1** | **#2** | **#3** | **#4** | **#5** | **#6** | **#7** | **#8** | **#9** | **#10** |
| ***Family*** | | | | | | | | | | |
| Rhodopsin-like GPCR superfamily signature | 9-30 | 54-76 | 90-111 | 139-162 | 292-316 | 329-255 |  |  |  |  |
| Muscarinic acetylcholine receptor family | 24-34 | 55-64 | 74-88 | 111-121 | 132-143 | 319-337 | 350-364 |  |  |  |
| G-protein coupled receptors family 1 signature | 60-76 |  |  |  |  |  |  |  |  |  |
| G protein coupled receptor, rhodopsin-like | 2-355 |  |  |  |  |  |  |  |  |  |
| ***Domain*** | | | | | | | | | | |
| GPCR, Rhodopsin like, 7TM | 1-347 |  |  |  |  |  |  |  |  |  |
| ***Unintegrated*** | | | | | | | | | | |
| Family A G protein-coupled receptor-like | 2-170 | 278-374 |  |  |  |  |  |  |  |  |
| 7tmA_mAChR | 1-358 |  |  |  |  |  |  |  |  |  |
| 5-hydroxytryptamine receptor | 2-367 |  |  |  |  |  |  |  |  |  |
| ***Sites*** | | | | | | | | | | |
| ligand binding domain | 54-55 | 58-59 | 106 | 139 | 142-143 | 146-147 | 307 | 310-311 | 333 | 336-337 |

A phylogenetic tree was made for the acetylcholine receptor with bootstrap support (SI Figure 9). The species and accession numbers used to construct the tree can be viewed in SI Table 15. Some nodes had lower bootstrap support, making it difficult to determine some placements within the tree. The *C. teleta* sequence clustered close to the *L. japonica* and *M. trossulus* sequences (SI Figure 9).

**SI Table 15: Accession numbers for the species and sequences used to make the acetylcholine receptor phylogenetic tree.** Accession numbers were found using NCBI blastp.

| **Acetylcholine Receptor** | **Accession** |
| --- | --- |
| ***Capitella teleta*** | ELT95769.1 |
| ***Homo sapiens*** | P11229.2 |
| ***Caenorhabditis elegan*** | NP_001024236.1 |
| ***Danio rerio*** | XP_001919160.1 |
| ***Xenopus laevis*** | XP_018115570.1 |
| ***Drosophila melanogaster*** | AAA28676.1 |
| ***Lytechinus pictus*** | XP_054758515.1 |
| ***Ciona intestinalis*** | XP_002121904.4 |
| ***Octopus sinensis*** | XP_029645888.1 |
| ***Liolophura japonica*** | XP_064600787.1 |
| ***Macrobrachium rosenbergii*** | XP_066977354.1 |
| ***Parasteatoda tepidariorum*** | XP_015911723.1 |
| ***Coccinella septempunctata*** | XP_044757106.1 |
| ***Mytilus trossulus*** | XP_063428510.1 |
| ***Anguilla anguilla*** | XP_035289578.1 |
| ***Oncorhynchus gorbuscha*** | XP_046154942.1 |
| ***Scyliorhinus canicula*** | XP_038663368.1 |
| ***Caretta caretta*** | XP_048710673.1 |
| ***Alligator sinensis*** | XP_006022579.1 |

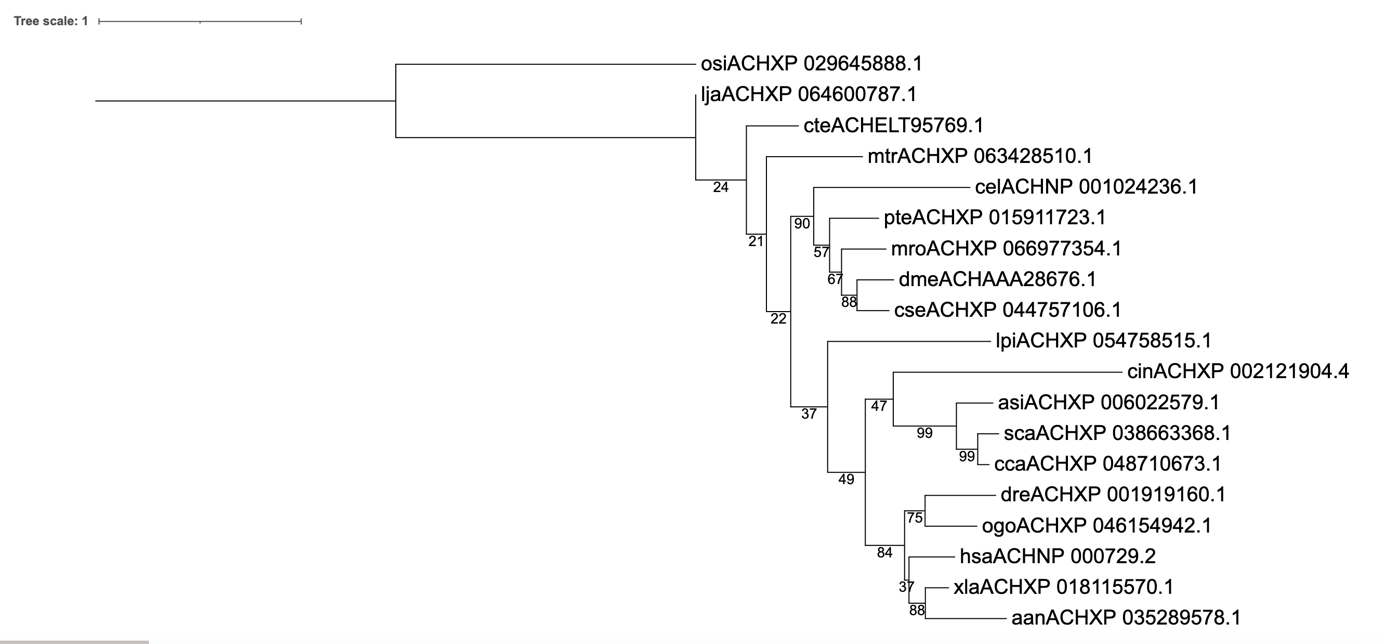

**Figure 9: Phylogenetic tree of acetylcholine receptor proteins across various species** Sequences were found using the *C. teleta* acetylcholine receptor sequence in NCBI blastp, then aligned using Clustal Omega. Alignments were masked using TrimAI and phylogenetic analyses were run using RaxML with 100 bootstraps. A LG4X+I model of amino acid substitution was used for the phylogenetic analyses. Nodes are labelled with the bootstrap support (%; 100 bootstraps). Species acronyms are: aan: *Anguilla anguilla,* Asi: *Alligator sinensis,* cel: *Caenorhabditis elegans,* cca: *Caretta caretta,* cin: *Ciona intestinalis*, cse: *Coccinella septempunctata,* cte: *Capitella teleta,* dme: *Drosophila melanogaster,* dre: *Danio rerio,* hsa: *Homo sapiens,* lja: *Liolophura japonica,* lpi: *Lytechinus pictus,* mro: *Macrobrachium rosenbergii,* mtr: *Mytilus trossulus,* ogo: *Oncorhynchus gorbuscha*, osi: *Octopus sinensis,* pte: *Parasteatoda tepidariorum,* sca: *Scyliorhinus canicular,* xle: *Xenopus laevis.*

GABA

Using the *C. elegans* and *H. sapiens* GABA-A receptor GABA-B receptor, 4-aminobutyrate aminotransferase, glutamate decarboxylase 1, and succinic semialdehyde dehydrogenase sequences, we found homologues in *C. teleta* (accession numbers can be found in SI Table 16). The specific values for these BLASTs can be found in SI Table 17. All *C. teleta* homologues were found using the *C. elegans* sequences. All BLASTs had low e-values, with the GABA-B Receptor and Glutamate Decarboxylase 1 having e-values of 0 (SI Table 17). All BLASTs had query covers 86% or higher and percent identities around 50 (SI Table 17). The GABA-A receptor had the lowest max score and percent identity of 298 and 37.47, respectively (SI Table 17). Once again, the human orthologues seemed to align to the *C. teleta* orthologues better than the *C. elegans* orthologues, having higher percent identities, positives, and lower gaps (SI Table 18).

**SI Table 16: Accessions for proteins involved in the GABA pathway from *Capitella teleta, Caenorhabditis elegans,* and *Homo sapiens****.* Accessions were taken off of NCBI

| **Protein** | **Capitella teleta** | **Caenorhabditis elegans** | **Homo sapiens** |
| --- | --- | --- | --- |
| GABA-A Receptor | ELU04795.1 | NP_499662.2 | P14867.3 |
| GABA-B Receptor | ELT91514.1 | NP_001256997.2 | Q9UBS5.1 |
| 4-aminobutyrate aminotransferase | ELT91408.1 | NP_501862.1 | P80404.3 |
| Glutamate Decarboxylase 1 | ELU00176.1 | NP_499689.1 | Q99259.1 |
| succinic semialdehyde dehydrogenase | ELU12107.1 | NP_001254393.1 | P51649.2 |

**SI Table 17: BLAST outputs from *Capitella teleta* for proteins involved in the GABA pathway.** BLASTS were run against *Caenorhabditis elegans* through NCBI blastp. Query Cover provides an indication of the length of each subject refers to the length of the target sequence compared to the reference sequence. Percent identity refers to the percentage of the amino acids that are identical between the sequences. Max score refers to the highest score calculated for the alignment between the sequences. The E-value refers to the number of alignments expected by chance with the calculated score.

| **Sequence** | **Query Cover (%)** | **Percent Identity** | **Max Score** | **E-value** |
| --- | --- | --- | --- | --- |
| GABA-A Receptor | 90 | 37.47 | 298 | 1e^-96^ |
| GABA-B Receptor | 87 | 44.18 | 641 | 0 |
| 4-aminobutyrate aminotransferase | 93 | 43.46 | 427 | 1e^-146^ |
| Glutamate Decarboxylase 1 | 86 | 57.99 | 550 | 0 |
| succinic semialdehyde dehydrogenase | 95 | 50.84 | 506 | 9e^-177^ |

**SI Table 18: Alignment information between *Capitella teleta* and *Caenorhabditis***

***elegans, and Capitella teleta* and *Homo sapiens* for proteins involved in the dopamine pathway.** Alignments were done using NCBI Global Alignments. Percent identity refers to the percentage of the amino acids that are identical between the sequences. Positives refers to the amount of amino acids that are the same or have similar properties between each sequence. Gaps refer to the amount of spaces in the alignment to account for any insertions/deletions in one of the sequences.

| **Species** | **Percent identity** | **Positives (%)** | **Gaps (%)** |
| --- | --- | --- | --- |
| **GABA-A Receptor** | | | |
| *C. teleta* vs. *C. elegans* | 37 | 66 | 6 |
| *C. teleta* vs. *H. sapiens* | 43 | 62 | 3 |
| **GABA-B Receptor** | | | |
| *C. teleta* vs. *C. elegans* | 44 | 62 | 3 |
| *C. teleta* vs. *H. sapiens* | 53 | 71 | 1 |
| **Choline Acetyltransferase** | | | |
| *C. teleta* vs. *C. elegans* | 43 | 63 | 0 |
| *C. teleta* vs. *H. sapiens* | 51 | 71 | 0 |
| **ATP citrate lyase** | | | |
| *C. teleta* vs. *C. elegans* | 58 | 74 | 0 |
| *C. teleta* vs. *H. sapiens* | 62 | 77 | 0 |
| **acyl-CoA synthetase short chain family member 1** | | | |
| *C. teleta* vs. *C. elegans* | 51 | 69 | 0 |
| *C. teleta* vs. *H. sapiens* | 60 | 79 | 0 |

The differential gene expression analyses showed that *C. teleta* contain all of the proteins examined in the GABA pathway. Succinic semialdehyde dehydrogenase had the highest expression, followed by 4-aminobutrate aminotransferase, GABA-B receptor, glutamate decarboxylase 1, and then GABA-A receptor (SI Figure 10).

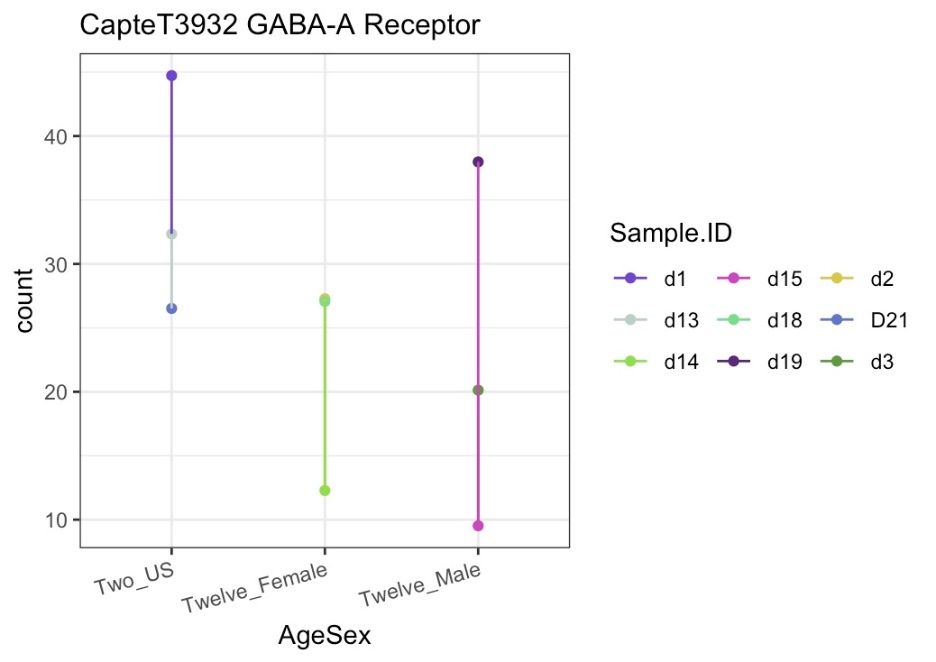

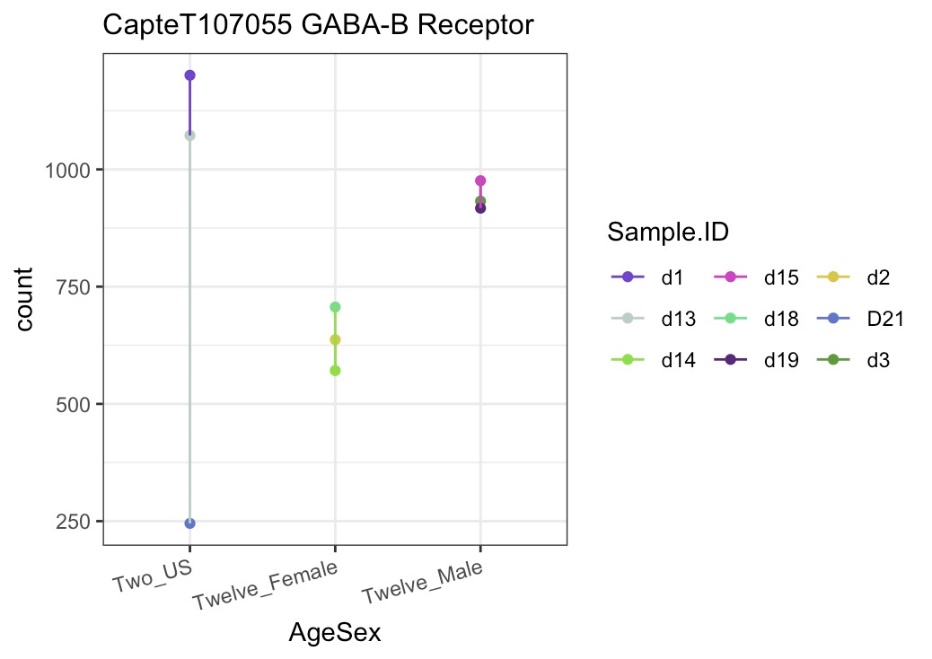

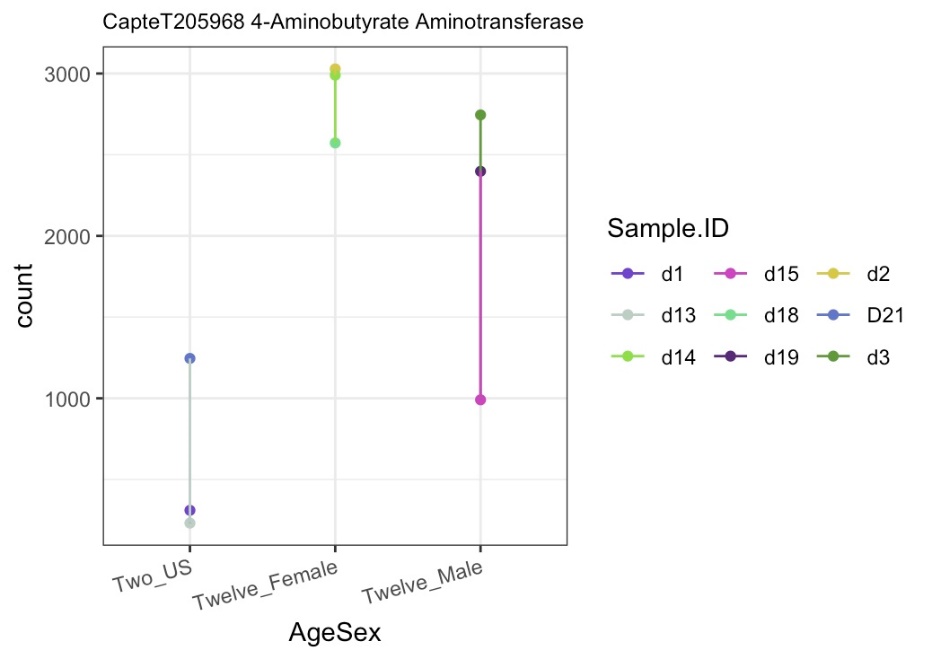

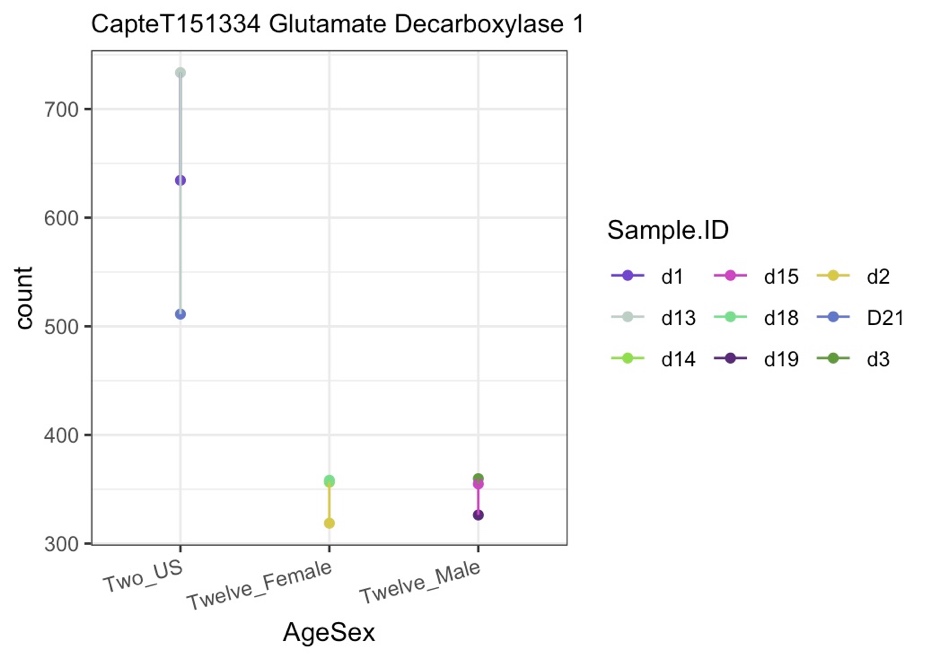

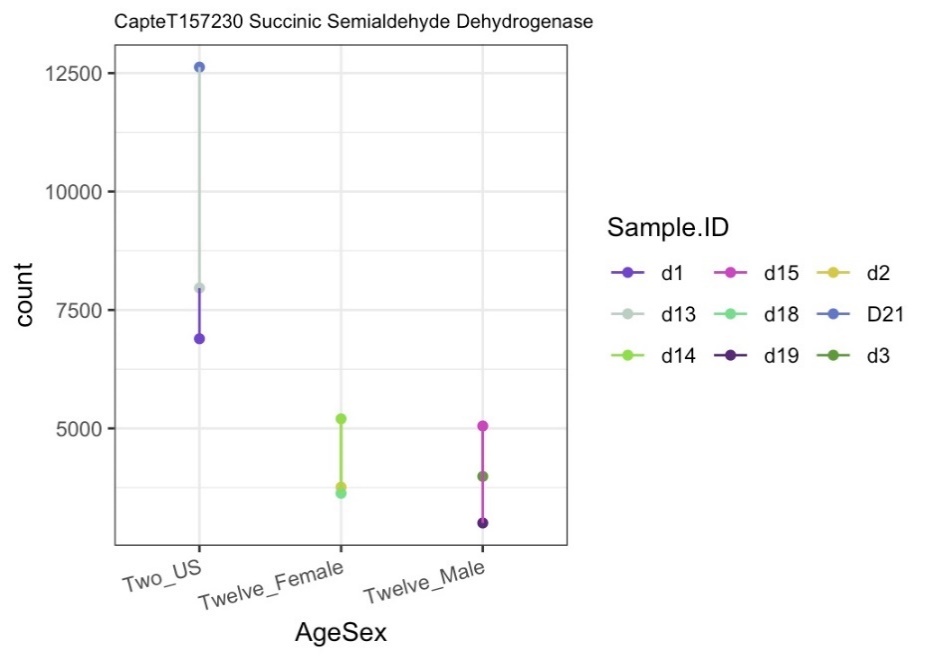

**SI Figure 10: Expression counts of proteins involved in the GABA pathway for juvenile, female, and male *Capitella teleta*.** The plots show the GABA-A receptor (A), GABA-B receptor (B), 4-aminobutrate aminotransferase (C), glutamate decarboxylase 1 (D), and succinic semialdehyde dehydrogenase (E). The different groups are Two_US (juveniles), Twelve_Female (females), and Twelve_Male (males). *C.* *teleta* juveniles were sampled at 2 weeks old and the males and females were sampled at 12 weeks old. There were 3 biological replicates for each group, each of a pool of 10 worms and 5 worms for juvenile and adult worms, respectively.

Using InterProScan with the *C. elegans,* *C. teleta,* and *H. sapiens* GABA-A receptor sequences, several motifs were found to be conserved between the species, including the gamma-aminobutyric-acid A receptor (alpha subunit), neurotransmitter-gated ion channel family signature, gamma-aminobutyric acid A receptor/Glycine receptor alpha, and All motifs were found *C. teleta* and are present in either/both the *C. elegans* and *H. sapiens* sequences in very similar locations within each gene (SI Figure 11, SI Table 19).

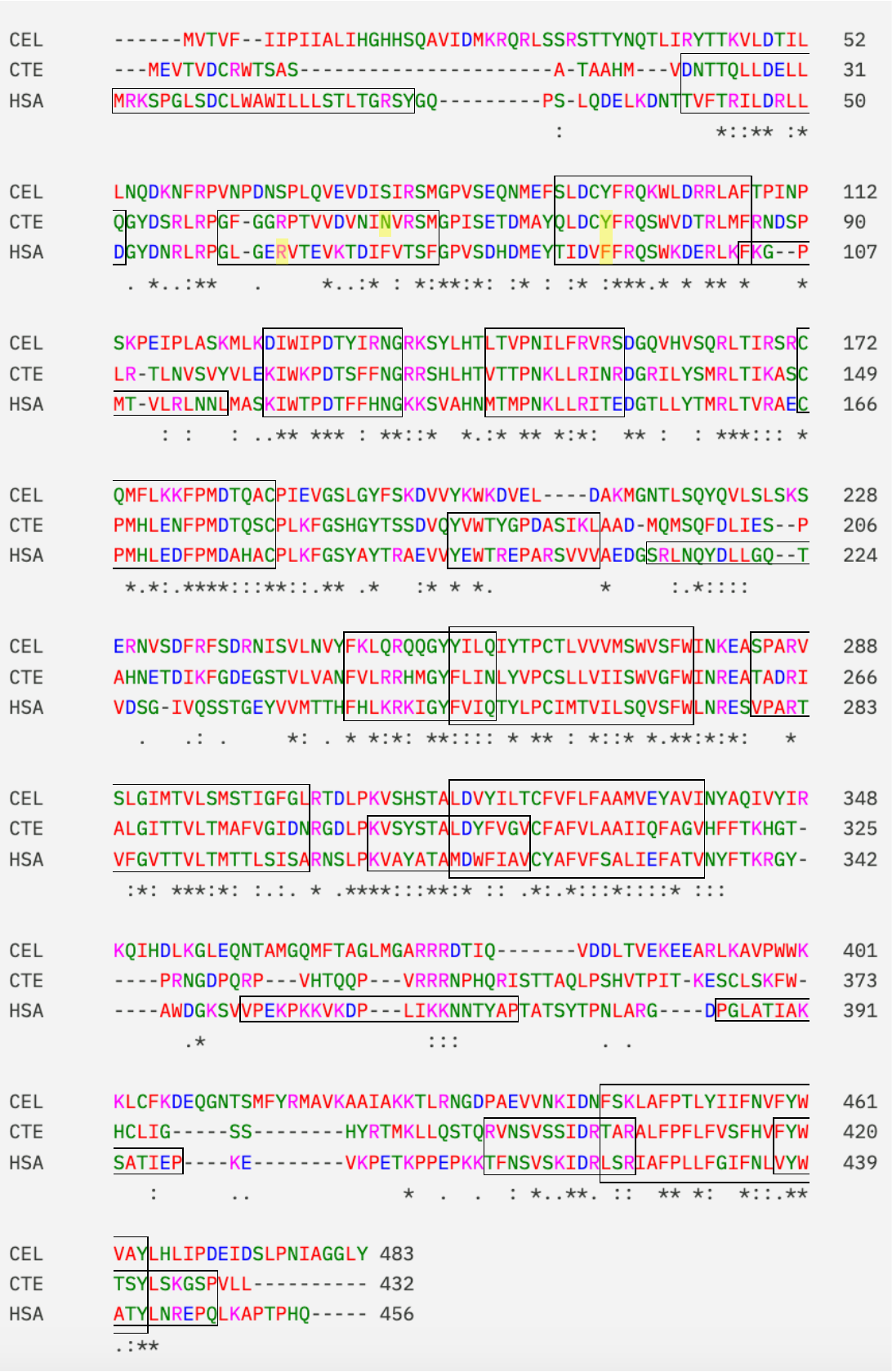

**SI Figure 11: Global alignment of acetylcholine receptor sequences between *C. teleta, H. sapiens,* and *C. elegans****.* Boxes indicate conserved areas between different sequences. The yellow highlights indicate ligand binding domains. * indicates positions that have a fully conserved residue. : indicates strong conservation between groups that have similar properties. . indicates weak conservation between groups that have similar properties.

**SI Table 19: Motif information for the *Capitella teleta* GABA-A receptor.** This information was found using InterProScan. Domains represent functional, structural, or sequence units. Family refers to groups of proteins that share common evolutionary origins, seen as similar functions, sequences, or protein structures. A homologous superfamily represents groups of proteins that share common evolutionary origins, reflected as similarity in the protein structure. Unintegrated represent motifs whose signatures are not classified. Sites refers to short sequences that have one or more conserved residues.

| **Amino Acid Numbers** | | | | | | |
| --- | --- | --- | --- | --- | --- | --- |
| **Motif** | **#1** | **#2** | **#3** | **#4** | **#5** | **#6** |
| ***Family*** | | | | | | |
| Neurotransmitter-gated ion-channel | 14-354 |  |  |  |  |  |
| Gamma-aminobutyric acid A receptor/Glycine receptor alpha | 236-256 | 262-283 | 296-317 | 403-423 |  |  |
| Neurotransmitter-gated ion channel family signature | 69-85 | 103-114 | 149-163 | 227-239 |  |  |
| Gamma-aminobutyric-acid A receptor, alpha subunit | 21-32 | 41-58 | 179-191 | 289-302 | 393-405 | 418-429 |
| ***Domain*** | | | | | | |
| LGIC_ECD_GABAR_GRD-like | 46-232 |  |  |  |  |  |
| LGIC_TM_anion | 235-420 |  |  |  |  |  |
| Neurotransmitter-gated ion-channel ligand-binding domain | 24-233 |  |  |  |  |  |
| Neurotransmitter-gated ion-channel transmembrane domain | 240-327 |  |  |  |  |  |
| ***Homologous Superfamily*** | | | | | | |
| Neurotransmitter-gated ion-channel ligand-binding domain superfamily | 11-232 |  |  |  |  |  |
| Neurotransmitter-gated ion-channel transmembrane domain superfamily | 230-428 |  |  |  |  |  |
| ***Residues*** | | | | | | |
| Ligand binding site | 54 | 73 |  |  |  |  |

A phylogenetic tree was made for the GABA-A receptor with bootstrap support (SI Figure 12). The species and accession numbers used to construct the tree can be viewed in SI Table 20. Some nodes had lower bootstrap support, making it difficult to determine some placements within the tree. The *C. teleta* sequence clustered close to the *L. japonica*, *O. sinensis*, and *M. trossulus* sequences (SI Figure 12).

**SI Table 20: Accession numbers for the species and sequences used to make the acetylcholine receptor phylogenetic tree.** Accession numbers were found using NCBI blastp.

| **Species** | **Accession** |
| --- | --- |
| ***Capitella teleta*** | ELU04795.1 |
| ***Homo sapiens*** | P14867.3 |
| ***Caenorhabditis elegans*** | NP_499662.2 |
| ***Danio rerio*** | XP_005166139.1 |
| ***Xenopus laevis*** | XP_018085048.1 |
| ***Drosophila simulans*** | XP_002077031.1 |
| ***lytechinus pictus*** | XP_054754594.1 |
| ***Ciona intestinalis*** | XP_026691627.1 |
| ***Octopus sinensis*** | XP_029639936.1 |
| ***Liolophura japonica*** | XP_064609709.1 |
| ***Macrobrachium rosenbergii*** | XP_066937073.1 |
| ***Parasteatoda tepidariorum*** | XP_015925423.1 |
| ***Coccinella septempunctata*** | XP_044749311.1 |
| ***Mytilus trossulus*** | XP_063415600.1 |
| ***Anguilla anguilla*** | XP_035266322.1 |
| ***oncorhynchus gorbuscha*** | XP_046165306.1 |
| ***scyliorhinus canicula*** | XP_038631651.1 |
| ***Caretta caretta*** | XP_048717731.1 |
| ***Alligator sinensis*** | XP_006021801.1 |

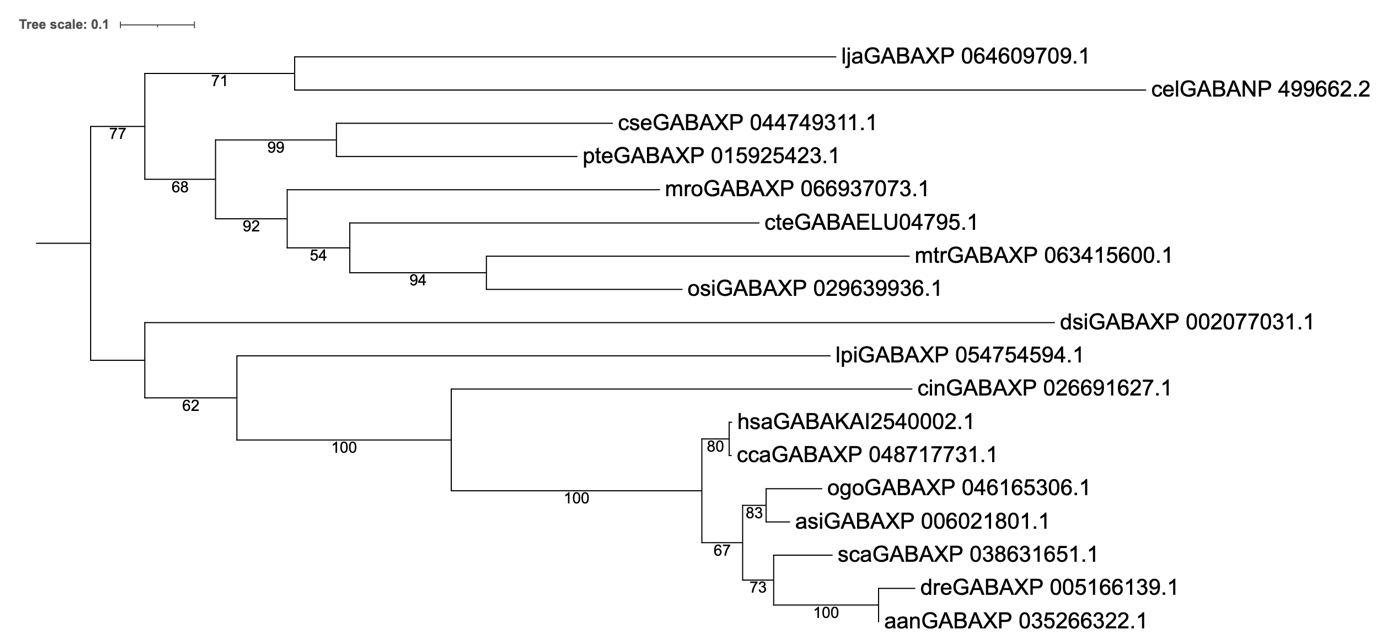

**Figure 9: Phylogenetic tree of GABA receptor proteins across various species.** Sequences were found using the *C. teleta* GABA receptor sequence in NCBI blastp, then aligned using Clustal Omega. Alignments were masked using TrimAI and phylogenetic analyses were run using RaxML with 100 bootstraps. A LC+4G model of amino acid substitution was used for the phylogenetic analyses. Nodes are labelled with the bootstrap support (%; 100 bootstraps). Species acronyms are: aan: *Anguilla anguilla,* asi: *Alligator sinensis,* cel: *Caenorhabditis elegans,* cca: *Caretta caretta,* cse: *Coccinella septempunctata,* cte: *Capitella teleta,* dme: *Drosophila melanogaster,* dre: *Danio rerio,* hsa: *Homo sapiens,* lja: *Liolophura japonica,* lpi: *Lytechinus pictus,* mro: *Macrobrachium rosenbergii,* mtr: *Mytilus trossulus,* ogo: *Oncorhynchus gorbuscha*, osi: *Octopus sinensis,* pte: *Parasteatoda tepidariorum,* sca: *Scyliorhinus canicular,* xle: *Xenopus laevis.*

Both the total distance moved (SI Figure 13) and average velocity (SI Figure 14) have peak and stable activity 1 hour after exposure to a novel environment. In the peak period, the average distance moved for juveniles in an equivalent 10-minute period would be 186.5 mm. The average velocity of the juveniles was 0.4 mm/s with a standard error of 0.1 mm/s in the peak and stable period of activity. Lastly, the average time to first movement in the juveniles was 270.2 s.

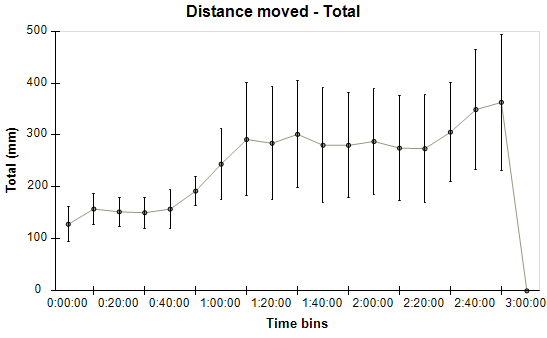

**SI Figure 12: Juvenile *C. teleta* (n=10) have increased distance moved after 1 hour in a novel arena.** Total distance (mm) was measured for 3 hours in 10-minute bins. Each point represents the mean total distance moved for each time bin. Error bars represent standard error of the mean. The graph does not go to the 3-hour mark because the software was not able to take measurements towards the end of the recording, resulting in only 1 *C. teleta* being measured.

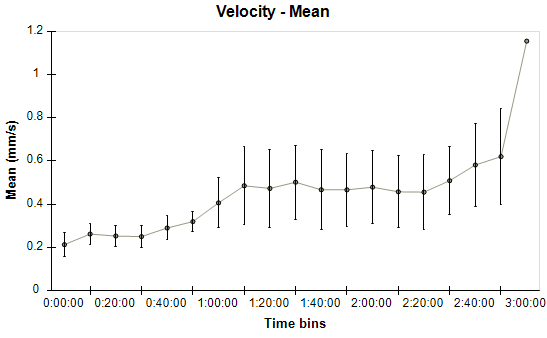

**SI Figure 13: Juvenile *C. teleta* (n=10) have an increase in average velocity 1 hour and 10 minutes after movement into a novel arena.** Velocity (mm/s) was measured for 3 hours in 10-minute bins. Each point represents the average velocity for each time bin. Error bars represent standard error of the mean. The graph does not go to the 3-hour mark because the software was not able to take measurements towards the end of the recording, resulting in only 1 *C. teleta* being measured.

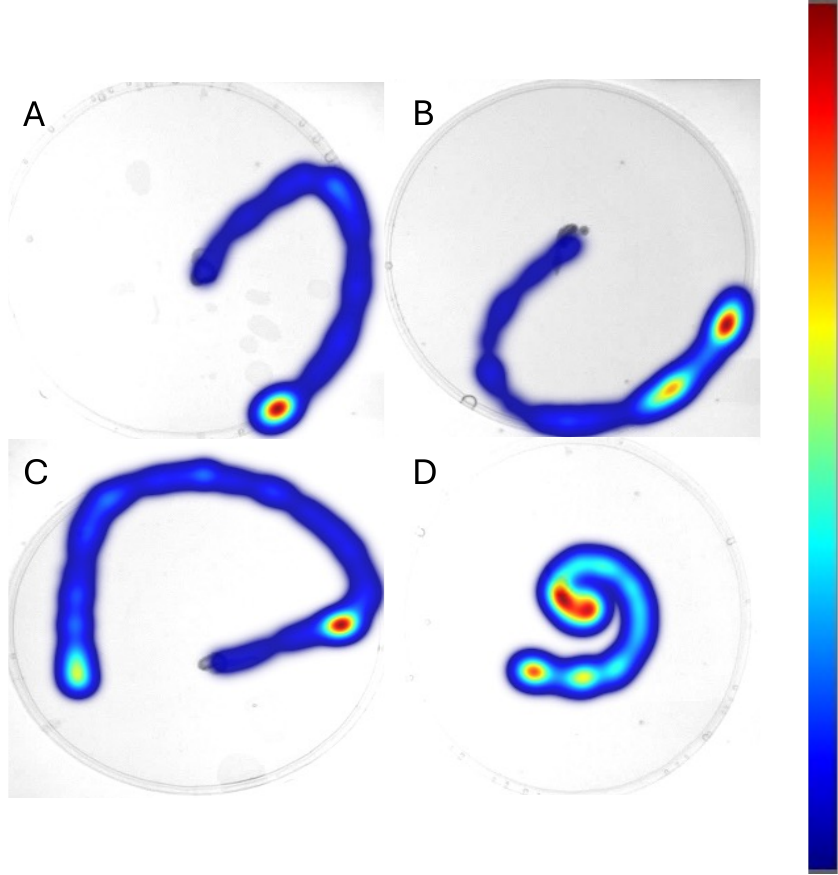

**SI Figure 14: Representative heat maps of adult *C. teleta* after exposure to nicotine in Petri dishes.** Adult *C. teleta* worms (n=11-14 per treatment) were exposed to a seawater control (A), 0.2 (B), 2 (C), or 20 (D) µM nicotine for 1 h in darkness at 15°C. After exposure, animals were moved to a novel arena (petri dish) to induce exploratory behaviour. Worms were video recorded in darkness at 15°C for 10 minutes. The traces show the animals movement around the arena, with blue representing less time spent, and red representing more time spent in an area.

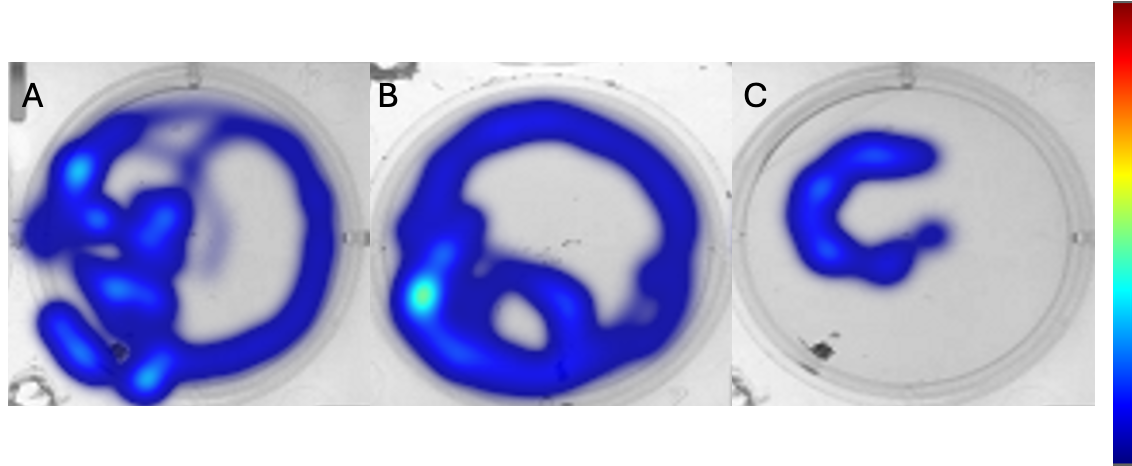

**SI Figure 15: Representative heat maps of adult *C. teleta* after exposure to nicotine in 6-well plates.** Adult *C. teleta* worms (n=12-14 per treatment) were exposed to a seawater control (A), 0.2 (B), 2 (C), or 20 (D) µM nicotine for 1 h in darkness at 15°C. After exposure, animals were video recorded in darkness at 15°C for 10 minutes. The traces show the animals movement around the arena, with blue representing less time spent, and red representing more time spent in an area.

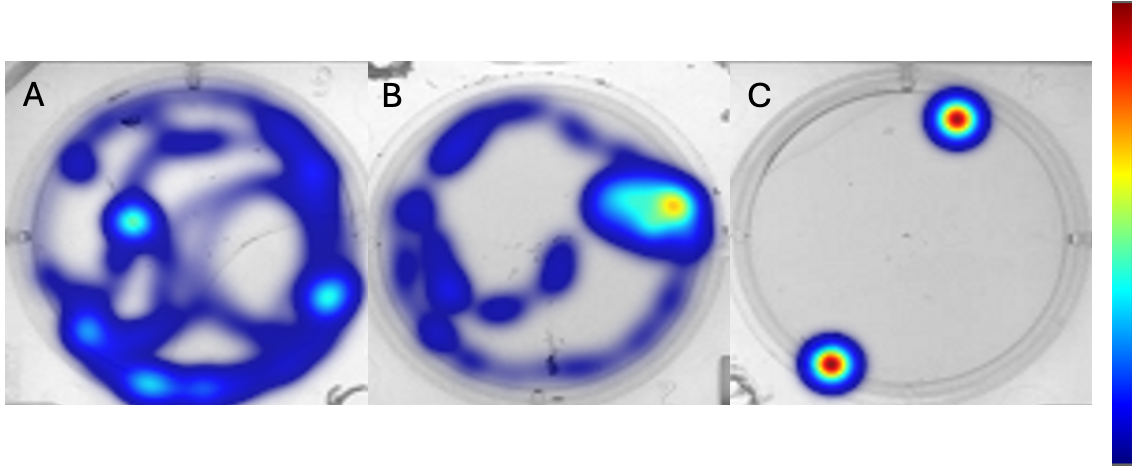

**SI Figure 16: Representative heat maps of juvenile *C. teleta* after exposure to nicotine in 6-well plates.** Adult *C. teleta* worms (n=12-13 per treatment) were exposed to a seawater control (A), 2 (B), or 20 (C) µM nicotine for 1 h in darkness at 15°C. After exposure, were video recorded in darkness at 15°C for 10 minutes. The traces show the animals movement around the arena, with blue representing less time spent, and red representing more time spent in an area.

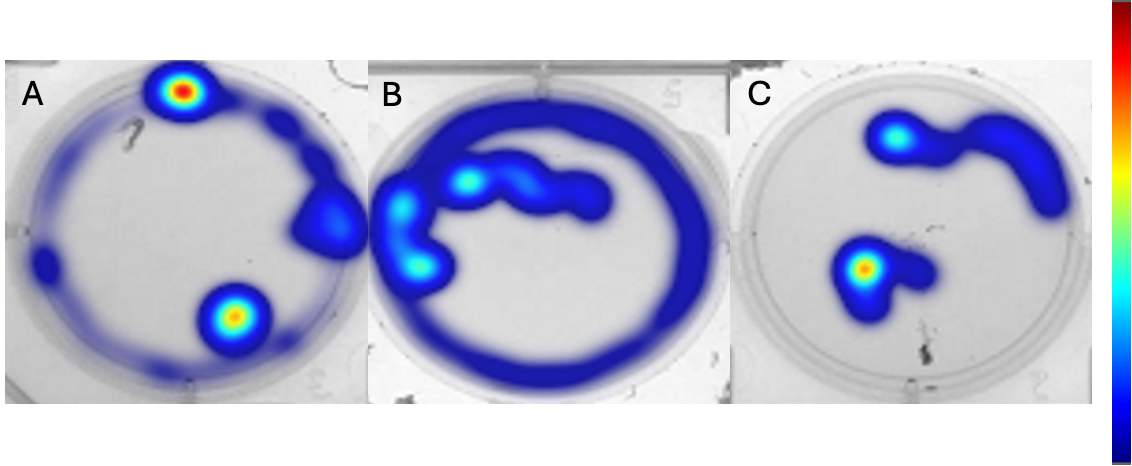

**SI Figure 17: Representative heat maps of juvenile *C. teleta* after exposure to fluoxetine in 6-well plates.** Adult *C. teleta* worms (n=12 per treatment) were exposed to a seawater control (A), 1 (B), or 10 (C) µM nicotine for 1 h in darkness at 15°C. After exposure, animals were video recorded in darkness at 15°C for 10 minutes. The traces show the animals movement around the arena, with blue representing less time spent, and red representing more time spent in an area.

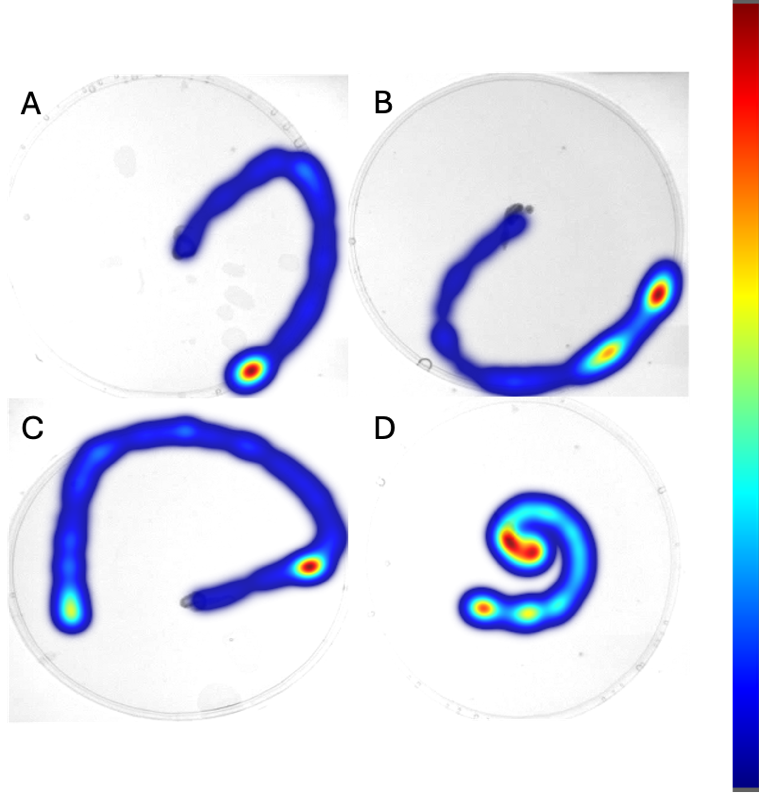

**SI Figure 18: Representative heat maps of juvenile *C. teleta* after exposure to apomorphine in 6-well plates.** Adult *C. teleta* worms (n=11-12 per treatment) were exposed to a seawater control (A), 0.01% DMSO (B), 1 (C), or 10 (D) µM nicotine for 1 h in darkness at 15°C. After exposure, animals were video recorded in darkness at 15°C for 10 minutes. The traces show the animals movement around the arena, with blue representing less time spent, and red representing more time spent in an area.

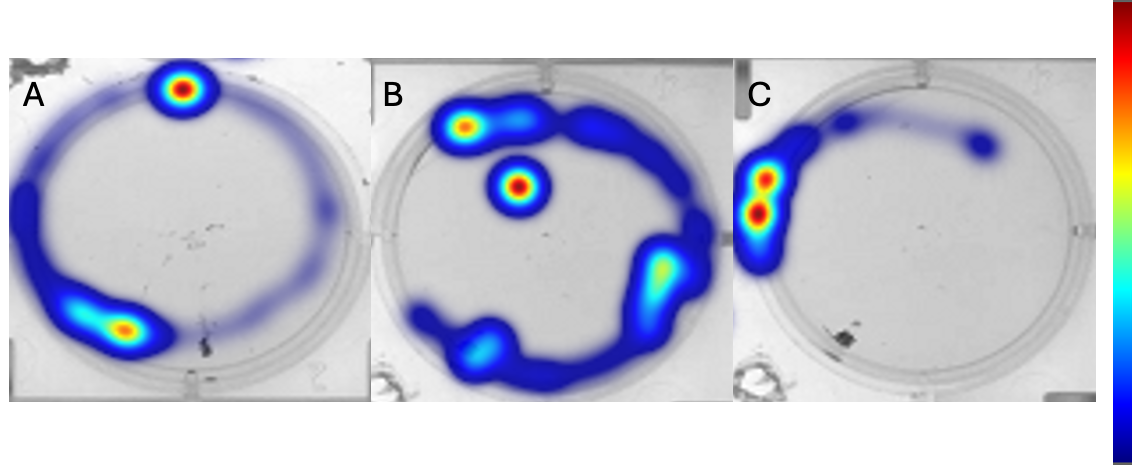

**SI Figure 19: Heat maps of juvenile *C. teleta* after exposure to phenobarbital in 6-well plates.** Adult *C. teleta* worms (n=11-14 per treatment) were exposed to a seawater control (A), 1 (B), or 10 (C) µM nicotine for 1 h in darkness at 15°C. After exposure, animals were video recorded in darkness at 15°C for 10 minutes. The traces show the animals movement around the arena, with blue representing less time spent, and red representing more time spent in an area.

***References***

D’Orbigny, A. (1835) ‘Histoire naturelle générale et particulière des Céphalopodes acétabulifères vivants et fossiles’, pp. 1–361.

Gould, A.A. (1850) ‘[descriptions of new species of shells from the United States Exploring Expedition]’, *Proceedings of the Boston Society of Natural History*, 3(151–156, 169–172, 214–218, 252–256, 275–278, 292–296, 309–312, 343–348), p. 344.

Pilsbry, H.A. (1893) ‘On Acanthopleura and its subgenera’, *The Nautilus*, 6(9), pp. 104–105.
